# A normative account of choice history effects in mice and humans

**DOI:** 10.1101/2020.07.22.216234

**Authors:** Junior Samuel Lopez-Yepez, Juliane Martin, Oliver Hulme, Duda Kvitsiani

## Abstract

Choice history effects describe how future choices depend on the history of past choices. Choice history effects are typically framed as a bias rather than an adaptive phenomenon because the phenomenon generally degrades reward rates in experimental tasks. However, in natural habitats, choices made in the past constrain choices that can be made in the future. For foraging animals, the probability of obtaining a reward in a given patch depends on the degree to which the animals have exploited the patch in the past. One problem with many experimental tasks that show choice history effects is that such tasks artificially decouple choice history from its consequences in regard to reward availability over time. To circumvent this, we used a variable interval (VI) reward schedule that reinstates a more natural contingency between past choices and future reward availability. By manipulating first- and second-order statistics of the environment, we dissociated choice history, reward history, and reaction times. We found that choice history effects reflect the growth rate of the reward probability of the unchosen option, reward history effects reflect environmental volatility, and reaction time reflects overall reward rate. By testing in mice and humans, we show that the same choice history effects can be generalized across species and that these effects are similar to those observed in optimal agents. Furthermore, we develop a new reinforcement learning model that explicitly incorporates choice history over multiple timescales into the decision process, and we examine its predictive adequacy in accounting for the associated behavioral data. We show that this new variant, known as the double trace model, has a higher predictive adequacy of choice data, in addition to better reward harvesting efficiency in simulated environments. Finally, we show that the choice history effects emerge in optimal models of foraging in habitats with diminishing returns, thus linking this phenomenon to a wider class of optimality models in behavioral ecology. These results suggests that choice history effects may be adaptive for natural contingencies between consumption and reward availability. This concept lends credence to a normative account of choice history effects that extends beyond its description as a bias.

## Introduction

Numerous perceptual and decision-making tasks have shown that choices systematically depend on the history of past choices, often referred to as choice history bias or choice inertia (Akrami, Kopec, Diamond, & Brody, 2018; Bari et al., 2019; Hwang, Dahlen, Mukundan, & Komiyama, 2017; Hwang et al., 2019; Fernberger, 1920; Fritsche, Mostert, & de Lange, 2017; Busse et al., 2011; Fründ, Wichmann, & Macke, 2014). This phenomenon is often framed as a bias rather than an adaptive phenomenon because it typically degrades the rate of reward consumption obtained by animals in experimental contexts. However, these studies predominantly make the implicit assumption that environmental resources are decoupled from the choices that animals make. In other words, such studies assume that past choices have no bearing on future reward availability. Such an assumption is nearly always violated by virtue of the causal structure of natural habitats. For instance, foraging for food in one patch may deplete the patch’s energy resources in the short term, whereas a patch left for longer may allow for an increased likelihood of replenishment. Thus, past choices impact the local foraging environment and affect outcome statistics. The variable interval (VI) reward schedule task is a simple task that circumvents this problem by making reward availability contingent on choice history. In the discrete version of the VI task, in each trial rewards are assigned with fixed or set reward probabilities keyed to different options. Such options could constitute the left or right port entry for mice or the left or right arrow key press on a computer keyboard for humans. Once assigned, the reward remains available until the animal chooses the associated option and consumes the reward. Once consumed, the reward is not available until it is assigned again to the same option (Zuriff, 1970). As a result, the reward probabilities in each trial differ from the set reward probabilities because they can change as a function of the animal’s past choices. In other words, the probability of consuming a reward, which is conditional on choosing a given option, is a function of the number of trials that have elapsed since the option was last chosen. Choosing the option with the highest set reward probability does not maximize the reward rate in this task, because even if the option has a lower set reward probability, the longer the time since that reward was last chosen the higher the chance that the reward will be assigned to a given option. Therefore, after a certain period, the probability that a reward is assigned to an unchosen option escalates until the option becomes that with the highest reward probability.

The best strategy for maximizing the reward rate over all of the available options thus depends not only on the set reward probability of each option but also on the history of how recently each option has been chosen in the past. To maximize their reward rate, optimal agents should choose the options with the highest set reward probability and should then switch when the reward probability of the other unchosen options overtakes that of the initial option. This strategy can resemble choosing the best option, with occasional switches to lesser options(Houston & McNamara, 1981). This choice strategy is consistent with the matching law(Herrnstein, 1961), which requires optimal agents to guide their decisions based on both past choices and set reward probabilities.

If a VI task structure mimics foraging in natural habitats, conditioning choices based on both past choices and past rewards may also emerge in optimality models that attempt to generalize to more ecological settings. Habitats in which foraging offers diminishing returns require optimal agents to follow the rules dictated by Marginal Value Theory(Charnov, 1976)] (MVT). MVT states that optimal agents should leave a food patch when the local reward rate from that patch decreases to a value that is below the average reward rate of the habitat (Bettinger & Grote, 2016). The same reward rate maximization that dictates optimal behavior in the VI task also dictates the behavior of an agent following MVT (Kubanek, 2017). Under MVT, the relative ratio of the reward rates between the chosen versus unchosen options determines the decision rule that is followed to switch options or, equivalently, to leave a patch. The apparent similarity between the natural contingencies studied by MVT and the contingincies explored artificially in VI tasks raises the following question: are similar behavioral strategies used by animals?

Here, we used a VI task to study foraging behavior from a normative perspective. We acquired behavioral data in both humans and mice and compared the choice history effects that we observed to those generated by optimal agents. By manipulating first- and second-order statistics of the reward outcomes, we characterized the contingency between choice and reward availability that may drive choice history effects. This result prompted us to derive a local decision rule by incorporating the choice history effects into a reinforcement learning algorithm. We call this RL variant the Double Trace (DT) model because it models choice history with both fast and slow decaying exponential functions. We find that in species as diverse as mice and humans as well as published data from monkeys (Lau & Glimcher, 2005), we can qualitatively observe the same choice history effects as those predicted from a normative perspective. Thus, we provide an explanatory account of choice history effects beyond their description as a bias. This finding potentially connects choice history phenomena to a broader class of optimality models within behavioral ecology.

## Results

Previous research in humans, monkeys, rats, and mice performing the two alternative free choice (2AFC) task showed that rewards reinforce choices toward choosing the same option and that this effect decays over time. This finding means that the further back in time a reward has occurred, the less influence that the reward exerts on the current choices (Lau & Glimcher, 2005; Kim, Sul, Huh, Lee, & Jung, 2009; Bari et al., 2019; Corrado, Sugrue, Seung, & Newsome, 2005). The dependency of choice on the reward history is sometimes referred to as a reward history effect (or reward bias). In contrast to reward history effects, past choices exert more complex effects on future choices. All else being equal, there is evidence that having chosen a given option recently, the probability of it being chosen again is decreased, in a phenomenon known as short-term alternation (Lau & Glimcher, 2005). This phenomenon coexists with long-term perseverance in which choice probabilities rebound, such that options chosen in the more distant past can again increase in their probability of being chosen again (Lau & Glimcher, 2005). Collectively, all of these types of effects are referred to as choice history effects or biases. While different species show similar reward history effects, choice history effects manifest with greater diversity. Namely, in most species, choice history effects show only a short-term alternation component and no long-term perseverance(Lau & Glimcher, 2005; Kim et al., 2009; Bari et al., 2019; Rutledge et al., 2009), with one exception being in non-human primates (Lau & Glimcher, 2005). This difference could be due to differences in task design or species-specific behavioral strategies. Furthermore, without normative models of behavior, it is difficult to argue why one form of choice history effect or another should exist. To address this problem, we test the performance of mice, humans, and optimal agents on a discrete version of the VI task with two alternative options (Lau & Glimcher, 2005). We also present the performance of monkeys in a VI task from a previously published work (Lau & Glimcher, 2005). This task is a variant of the 2AFC task with a VI schedule of reinforcements. Henceforth, we refer to this task simply as the VI task. In this task, the water rewards of two alternative options are delivered probabilistically following a baiting schedule (Fig. 1b, eq. 1). First, we describe the choices of a reward maximizing agent. To do so, we construct an optimal agent that has full knowledge of the objective reward probabilities and task structure. Hence, we refer to this agent simply as the Oracle agent. We also construct an optimal agent that uses Bayesian inference to optimally infer the reward probabilities based on prior outcome experiences. Note that this agent updates its beliefs regarding the unchosen option following the VI schedule of reinforcements (eq. 1). We refer to this agent simply as the Bayesian agent. The optimal agents are described fully in the Methods section for optimal agents. In the following, we use linear regression analysis to describe what behavioral contingencies drive choices in mice and humans, and then, we formally compare this analysis to the behavior of the optimal agents.

**Figure 1:**
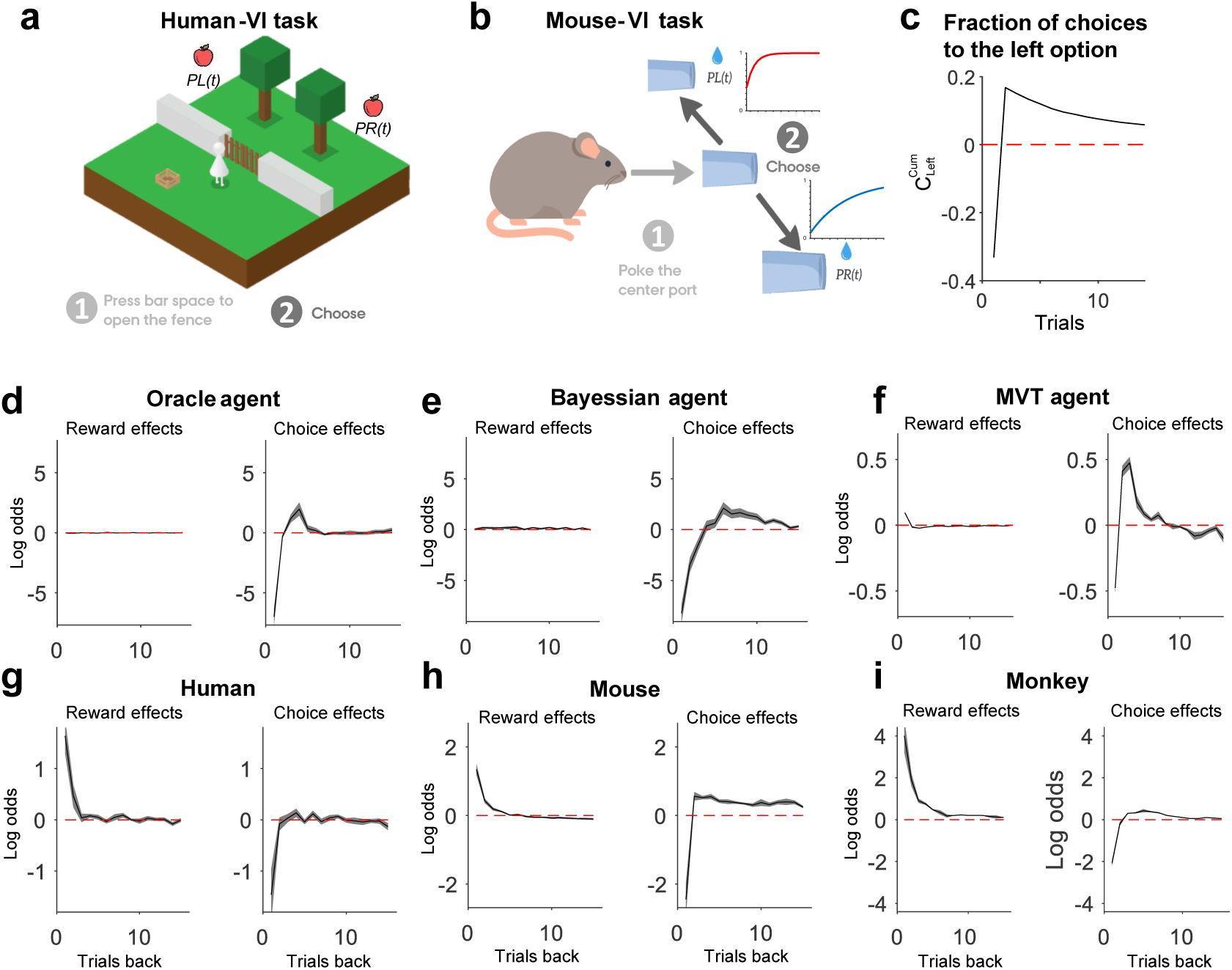
Reward and choice history effects in animals, humans and optimal agents. (a) Snapshot of the computer game played by the human participants. The subjects had to wait between 0s and 5s after opening a virtual fence by pressing on a keyboard before making the decision to press on the left or right key. (b) The scheme of the task was adapted for mice. The rodents had to poke the center port to start a trial and wait in the center port 0.2-0.4s before choosing the right or left port. For (a) and (b), the reward is assigned probabilistically to the alternative options independently of whether or not an agent visits the option in the given trial, and it remains to be collected until an agent chooses that option. In other words, rewards are allocated under a baiting schedule. The baiting schedule results in an increased probability of obtaining a rewards from an unchosen options, as illustrated in (b) with red and blue curves when reward delivery probabilities are 0.4 and 0.1 respectively. The baiting scheduel is described by eq.1 (c) shows the ideal fraction of choices allocated to the same options in a two-alternative task under the baiting schedule where the set reward probabilities of each option are taken randomly between 0 and 1 from a uniform distribution. (d-i) The influence of past rewards and choices on the current choice. (d), the Oracle agent (10 sessions) (e), the Bayesian agent (10 sessions) (f) and the MVT agent (25 sessions), (g) humans (19 sessions and 19 subjects) (h), rodents (219 sessions and 7 mice) was analysed using logistic regression with elastic-net regularisation. The coefficients with the lowest deviance in a five-fold cross-validation process were selected. For the MVT agent in (f), the maximum number of rewards in a patch was *Gmax* = 100, and the maximum travel time was *travelmax* = 108.33 s. The depletion rate of each patch followed an exponential distribution with a mean of 0.25. The bold lines depict the average, and the shaded regions depict the standard error of the mean (s.e.m.). (i) Lag regression coefficients indicating the influence of recent rewards (left) and choices (right) on the current choices of two monkeys from Lau and Glimcher’s study(Lau & Glimcher, 2005).

### Optimal agents show short-term alternation and long-term perseverance behaviors

The reward probabilities in the VI task that we used to test all agents (mice, humans, and synthetic) were updated (baited) in each trial according to the following rule:

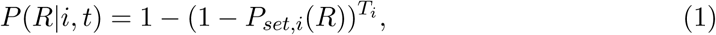

where *P*(*R*|*i, t*) represents the probability of receiving reward *R* at trial *t* for action *i*. This probability depends both on the number of trials *T*_*i*_ since the last time that the same action *i* was chosen and on the set probability *P*_*set,i*_(*R*) of scheduling a reward for the option *i* set by the experimenter.

First, we ran optimal agents over multiple sessions in the VI task. For each trial (n = 200,000 trials in total), set reward probabilities were drawn from a uniform distribution *P*(*R*|*l*) ∈ [0 1], *P*(*R*|*r*) ∈ [0 1] for left (*l*)- and right (*r*)-sided options. The uniform distribution on the interval of [*a b*] was henceforth defined as 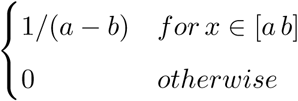. Next, we quantified how much the initial reward probabilities favored the choice for the same versus the alternative option on trial *t* given that the probabilities (updated according to eq. 1) in all previous trials favored the choices for the same option. Here, we consider past choices only for the left side, although the same results are obtained for the right-hand choices. Choices were defined to take values of 1 and –1 values for the left-hand and right-hand sides, respectively (following eq. 2):

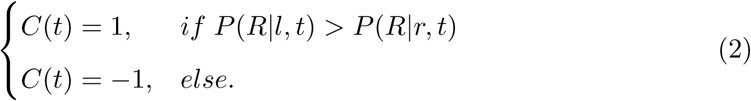

We computed the sum of the differences between the left and right choices given that the probabilities favored the left choices in all previous trials, following the equation:

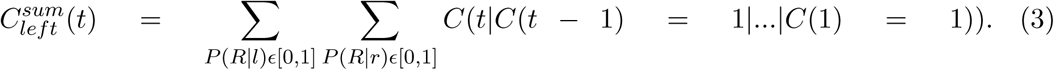

Here, 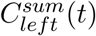 is the cummulative number of left choices minus the cumulative number of right choices on a current trial *t* provided that the probabilities favored the left choices in all previous trials. We observed that short-term alternation was favored over long-term perseverance for the second trial. All subsequent trials favored perseverence over alternation (Fig. 1c). This analysis revealed how the probabilities in the VI task favored alternation over perseverence as a function of previous choices. Another way to quantify the effect of past choices and rewards on future choices is to use the agent’s past rewards and past choices to predict its future choices. To do so, we analyzed the choice dynamics using logistic regression. The Oracle agent was an oracle insofar as it “knew” in advance the reward probabilities of each option on each trial. Each session consisted of 500 trials with set reward probabilities drawn from a uniform probability distribution *P*(*R*|*l*) ∈ [0 1], *P*(*R*|*r*) ∈ [0 1] for left (*l*) and right (*r*) sided options. We generated 100 such sessions and used regularized regression analysis with a cross-validated elastic net (Methods, see Linear regression) (Hastie, Tibshirani, & Friedman, 2009) to reveal how the past choices and rewards affected the future choices. This analysis revealed strong alternation effects for immediate past choices, and perseverance for choices further back into the past (Fig. 1d). The beliefs of the Bayesian agent were characterized by a flat prior distribution over reward probabilities *P*(*R*|*i*) ∈ [0 1] for both the left and right sides, and the agent updated these beliefs for the chosen options according to its experience. For the unchosen option, the agent updated the probabilities according to the baiting schedule (eq. 1, Methods section on optimal agents). The Bayesian agent was run for 1000 separate sessions with set reward probabilities drawn from a flat uniform distribution ranging from *P*(*R*|*i*) ∈ [0 1] for both the left- and right-hand sides. Each session consisted of 1000 trials. The choice and history effects of the Bayesian agent were estimated by the same regression analysis as for the Oracle agent. Both the Oracle and Bayesian agents generated similar choice history effects that indicated strong short-term alternation and mild long-term perseverance (Fig. 1c-e, right panels). It should be noted that by virtue of its knowledge of the objective probabilities, the past rewards did not influence the choices of the Oracle agent. This was not true for the Bayesian agent, which was most influenced by reward outcomes at the begining of the session. As these probabilities are inferred with greater certainty (as experience accrues), the agent’s choices are increasingly guided by the same probabilities (Fig. 1c-e, left panels). In other words, the performance of the Bayesian agent approached the performance of the Oracle agent toward the end of the sessions, and likewise, the Bayesian agent became less sensitive to past rewards.

### Short-term alternation and long-term perseverance of choices in optimal agents foraging in habitats not limited to two options

The VI task structure has traditionally been used to examine the behavior of animals faced with two options. However, foraging animals in natural habitats are typically faced with a far greater number of options.. Thus, the algorithms extracted from animal behavior in VI habitats might not necessarily capture the more general rules that drive animals’ choices. How should an optimal agent maximize its reward rate when faced with a large number of options? In such habitats, MVT formalizes a simple decision rule: leave a patch when the local reward rate from that patch drops below the average reward rate of the habitat (Charnov, 1976; Bettinger & Grote, 2016). In essence, an MVT agent compares two values that are in principle independent of the number of available options: the reward rate of the current patch, and the reward rate averaged globally over all relevent patches. To test what type of reward and choice history effects an MVT agent generates, we simulated habitats with heterogeneous patches with diminishing returns due to depletable resources (1000 virtual food patches). The patches differed in terms of their reward density, denoted as G, which was drawn from a uniform probability distribution within the range of [0, *G*_*max*_ = 100]. The depletion rate was drawn from an exponential distribution with a mean of 0.25, and the travel time was drawn from a uniform distribution of range [50, 108.33] seconds. We looked at the reward and choice history effects that could influence behavior. To our surprise, we observed that MVT agents generated both a short-term leaving strategy (analogous to alternation in the VI task) as well as a long-term staying strategy (analagous to perseverance in the VI task) strategy (Fig. 1f, right panel). The reward history effects reflect the tendency of such agents to stay on patches where they have recently been rewarded (Fig. 1f, left panel). The different travel times and depletion rate did not have observable effects on the MVT agent’s choices (Suppl. Fig. 2).

The following intuition may explain the observed short-term alternation and long-term perseverance of the choice effects of MVT agents. Due to the heterogeneity of the habitats, patches with different reward densities undergo different rates of depletion. The foraging agent’s decision whether to stay or to leave the patch is controlled by the depletion rate of rewards from the chosen patch on consecutive trials. On average, if the forager decides to stay in the same patch and collect rewards, the reward density will be lower compared to the average reward density of the global habitat. Because of this, the optimal strategy is to leave most of the patches after depleting their respective reward densities. However, the remaining patches that had either a higher density of initial rewards or a slower depletion rate that favored decisions to stay on a second trial will also favor staying decisons for subsequent trials. Thus, both alternation and perseverance, albeit at different temporal scales, determine the choice dynamics of the MVT agent. This is qualitatively similar to choice history effects that Oracle and Bayesian agents generate in VI task.

### Short-term alternation and long-term perseverance of choices in mice and humans

Next, we sought to elucidate whether the same choice history effects emerge in mice and humans performing the VI task. In contrast to the optimal agents, mice and humans can estimate poor versus rich options based on their reward history. In addition to this fact, we anticipated that our test subjects may need to keep track of the history of their choices, because the probability of reward for the unchosen option continually increases with time (eq. 1). Thus, it was expected that both reward and choice histories would contribute to future choices. To test this hypothesis, we ran both mice and humans on a VI task. Subjects in the VI task had to choose left- or right-sided options after initiating the trial with the center port nose poke in (in the case of mice) or pressing the space bar (in the case of human subjects). The specific set reward probabilities for the left-versus right-hand sides, respectively, that we used for the VI task were 0.25:0.25, 0.4:0.1, and 0.1:0.4 for mice (n = 7 animals) and 0.4:0.1 and 0.1:0.4 for humans (n = 19 subjects). Each block consisted of 50-150 trials for mice and 20-30 trials for humans. Each mouse completed 1-27 blocks of sessions, while human subjects completed at least 6 blocks per session. We also looked at published data from monkeys (non-human primates) (Lau & Glimcher, 2005). For the monkey data, the set reward probabilities for left-versus right-hand sides according to (Lau & Glimcher, 2005) were 0.23:0.075, 0.075:0.23 and 0.043:0.25, 0.25:0.043.

Logistic regression with elastic-net regularization (see Methods, linear regression) showed the monotonic diminishing effects of past rewards on upcoming choices. The past choices showed a mixture of short-term alternation and long-term perseverance (Fig. 1g,h). Logistic regression analysis of the monkey’s choice data also showed qualitatively similar effects of past rewards and choices on future choices (Fig. 1i). While choice history effects showed some diversity among species and optimal agents, one commonality among the species was that recently choosing a given option decreased its probability of being chosen again in the next trial, whereas choices made two or more trials back in history increased the probability of the same option being chosen again in the next trial.

### Dissociating reaction time, choice, and reward history effects

Our previous analyses showed that reward history and choice history exert separable effects on choice. First, we found that regression coefficients for rewards (mean(SD) = 1.34(0.42)) and choices (mean(SD) = -2.4(1.29)) were significantly different (p = 0.0006, Mann-Whitney U test, n = 7 animals) for the mice and human data (rewards: mean(SD) = 1.39(1.37); choices: mean(SD) = -1.17(1.93), p = 0.00001, Mann-Whitney U test, n = 19 human subjects). From this finding, it was not clear whether these effects reflected the same or different computational processes. To interrogate what drives choice and reward history effects, we manipulated the reward outcome statistics over three different dimensions. Specifically, we manipulated one of three behavioral contingencies: experienced total-reward rate, difference in reward delivery probabilities and volatility, while holding the other two contingencies constant. Besides reward and choice history effects, we also included reaction time (RT) as a metric of an animal’s performance. We defined RT as the time spent by the animal between leaving the center port to poking a side port. In other words, RT was the time spent by the animal taking an action after starting a trial. It has been reported that RTs can be driven by reward expectations(Lauwereyns, Watanabe, Coe, & Hikosaka, 2002; Zariwala, Kepecs, Uchida, Hirokawa, & Mainen, 2013), while reward history effects are sensitive to the volatility of reward outcomes (Behrens, Woolrich, Walton, & Rushworth, 2007). However, very little is known regarding what drives choice history effects. Here, we hypothesized that choice history effects reflect the growth of the reward probabilities of the unchosen options. The faster the reward probability of the unchosen option outgrows the reward probability of the chosen option, the stronger the alternation bias should be, and thus the first regression coefficient of the choice effects should be negative.

The effects of volatility on behavioral performance was analyzed by pooling the set of behavioral sessions in which block size was manipulated (4 animals, 44 sessions) and dividing the sets into three groups (terciles) of block lengths of either 14 - 54, 55 - 96, or 97 - 136 trials per block in a session. Our results showed that the volatility of the reward outcomes affected the reward history effects. The strongest effects were observed on the first regression coefficient for past rewards (1^st^tercile mean(SD) = 1.86(1.59), 2^nd^tercile mean(SD) = 1.24(0.89), 3^rd^tercile mean(SD) = 0.34(0.38), Pearson correlation coefficient (r) = -0.35, p = 0.0032, 95% CI [-0.54 -0.12]), while the regression coefficients for past immediate choices (1^st^tercile mean(SD) = -3.66(1.9), 2^nd^tercile mean(SD) = -2.49(1.88), 3^rd^tercile mean(SD) = .2.95(1.47), r = 0.24, p = 0.043, 95% CI [0.01 0.45]) and RTs (1^st^tercile mean(SD) = 0.19(0.04), 2^nd^tercile mean(SD) = 0.2(0.05), 3^rd^tercile mean(SD) = 0.24(0.04), r = 0.29, p = 0.016, 95% CI [0.06 0.49]) were less affected (Fig. 2a). Different from mice we did not observe any dependency between environmental volatility and the behavior of the Oracle agent (Suppl. Fig. 3b). As mentioned previously, this is due to the fact that the Oracle agent is not required to integrate its reward history into future choices because it already has access to the objective reward probabilities of each option.

**Figure 2:**
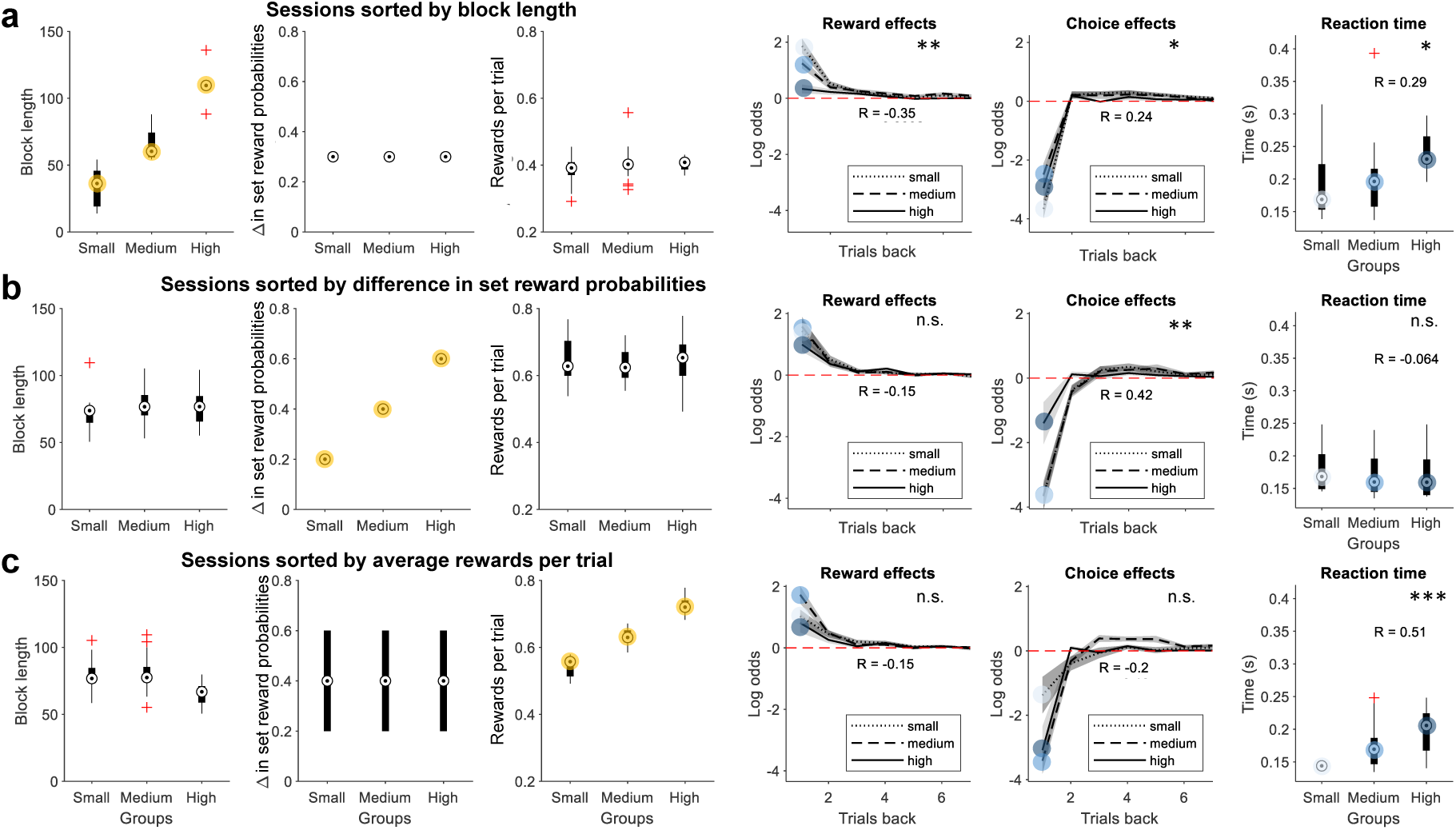
The effects of volatility, difference in the set reward probabilities, and reward rate per trial on reward and choice history effects and reaction times. (a) Sessions sorted into three groups, or terciles, based on the number of trials per block condition (4 mice in 44 sessions). The set reward probabilities pair was 0.1 vs. 0.4. The three left-hand panels show the distribution of the block lengths, the difference in the set reward probabilities, and the average number of rewards per trial obtained by the mice, in that order. The three right-hand panels show their corresponding reward and choice history effects, as well as the reaction times (the time from which the animal left the center port and poked a side port). (b) Sessions sorted into three groups based on the difference (Δ) in set reward probabilities (4 mice in 48 sessions). The set reward probabilities pair per session were 0.4 vs. 0.6, 0.3 vs. 0.7, and 0.2 vs. 0.8. The three left-hand panels show the distribution of the block lengths, the difference in the set reward probabilities, and the average number of rewards per trial (reward rate) obtained by the mice, in that order. The three right-hand panels show their corresponding reward and choice history effects, as well as the reaction times. (c) Sessions sorted in three groups based on the average number of rewards collected per trial (4 mice in 48 sessions). The set reward probabilities pair per session were 0.4 vs. 0.6, 0.3 vs. 0.7, and 0.2 vs. 0.8. The three left-hand panels show the distribution of the block lengths, the difference in the set reward probabilities, and the average number of rewards per trial obtained by the mice, in that order. The three right-hand panels show their corresponding reward and choice history effects, as well as the reaction times. The correlation of the block lengths with the regression coefficients of reward and choices one trial back, as well as with the reaction times, is reported as r on the plots with their corresponding significance labeled as *p* < 0.05 = *, *p* < 0.01 = **, *p* < 0.001 = * * *, and n.s. states for a non-significant result.

The effects of the difference in the set reward probabilities (0.8 vs. 0.2, 0.7 vs. 0.3, 0.6 vs. 0.4) on mice performance were analyzed by running animals on a different set of sessions (n = 4 mice, n = 48 sessions). Once more, we partitioned behavioral sessions into terciles based on the difference in the set reward probabilities. The choice history effects only analyzed for the first regression coefficients of the mice showed significant changes in response to the difference in the set reward probabilities (1^st^tercile mean(SD) = -3.66(1.48), 2^nd^tercile mean(SD) = -3.64(1.78), 3^rd^tercile mean(SD) = -1.4(2.56), r = 0.42, p = 0.0032, 95% CI [0.15 0.62]), while both reward history effects only analyzed for the first regression coefficients (r = -0.15, p = 0.32, 95% CI [-0.41 0.15]) and RTs (r = -0.064 and p = 0.66, 95% CI [-0.34 0.22]) did not show significant changes (Fig. 2b). These effects were also the same for the Oracle agent (Suppl. Fig. 3a).

**Table 1:**
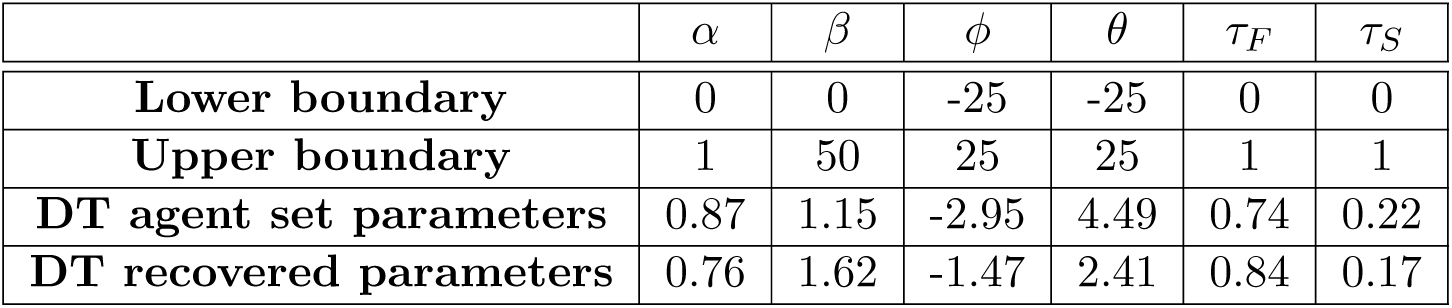
Recovered parameters during behavioural prediction from the choices and rewards of a DT agent running a virtual VI task.

| | $\alpha$ | $\beta$ | $\phi$ | $\theta$ | $\tau_F$ | $\tau_S$ |
| --- | --- | --- | --- | --- | --- | --- |
| **Lower boundary** | 0 | 0 | -25 | -25 | 0 | 0 |
| **Upper boundary** | 1 | 50 | 25 | 25 | 1 | 1 |
| **DT agent set parameters** | 0.87 | 1.15 | -2.95 | 4.49 | 0.74 | 0.22 |
| **DT recovered parameters** | 0.76 | 1.62 | -1.47 | 2.41 | 0.84 | 0.17 |

Finally, we partitioned the same set of sessions (n = 4 animals, n = 48 sessions) used in the previous analysis into terciles based on the reward rate experienced by the animal per trial. We found that RTs changed as a function of the experienced reward rates. The RTs showed a significant and strong positive correlation (1^st^tercile mean(SD) = 0.14(0.01), 2^nd^tercile mean(SD) = 0.17(0.03), 3^rd^tercile mean(SD) = 0.2(0.03), r = 0.51, p = 0.0002, 95% CI [0.26 0.69]) to the overall experienced reward rates (Fig. 2c, right-most panel), while reward (r = -0.15, p = 0.3, 95% CI [-0.42 0.14]) and choice history (r = -0.2, p = 0.18, 95% CI [-0.45 0.09]) effects were not significantly affected (Fig. 2c). These observed effects were counter-intuitive, because most other studies on this topic have shown a negative correlation between RTs and experienced reward rates (Lauwereyns et al., 2002; Zariwala et al., 2013). To understand this discrepancy, we must emphasize that the combined (left-vs. right-hand options) set reward probabilities per session were the same across conditions (for probabilities 0.2, 0.3, 0.4, 0.6, 0.7, and 0.8). Therefore, an increase in the number of rewards per trial experienced by the animals can only be due to the animal’s choice strategy. The fundamental structure of the VI task allows the reward rates (rewards per trial) to depend on both the animal’s choices as well as the reward delivery probabilities. We independently validated the negative correlations between the RTs and experienced reward rates by manipulating both the overall reward rates and set reward probabilities, as seen in Supplementary Figure 4.

### Double Trace model incorporates choice history effects into action selection

To incorporate the choice history effects into a local decision rule, we derived an RL model based on logistic regression (see Appendix for derivation of the DT model). The DT model captures the choice history effects by including the sum of two scaled choice traces *F* and *S*, hence the name ‘double-trace’ model (DT). These traces were updated with independent learning rates (eqs. 6 and 7). Reward history effects were captured using the value update equation with forgetting Q (Ito & Doya, 2009) (eq. 5). According to the DT model, the probability that an agent chooses action *i* (i.e., the right-or left-hand action) at time step *t* follows the softmax action selection rule:

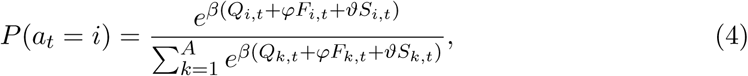

where *Q*_*i,t*_ denotes reward expectation, *F*_*i,t*_ is the fast choice trace, *S*_*i,t*_ is the slow choice trace, and *β* is the inverse temperature. The higher the inverse temperature, the more stochastic the choices are, and thus, this quantity can be thought of as an exploration factor. The scaling factors for the choice traces are *φ* and *ϑ*. Each choice trace has a learning rate of either *τ*_*F*_ or *τ*_*S*_. *F* and *S* choice traces extracted from animals typically decay at fast and slow rates, respectively (i.e., *τ*_*F*_ ≥ 0.5 and *τ*_*S*_ < 0.5).

The reward expectation and choice traces are updated with the learning rates of *α, τ*_*F*_, and *τ*_*S*_ as follows:

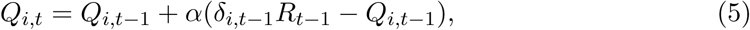

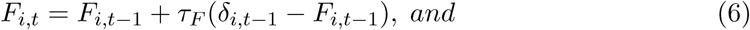

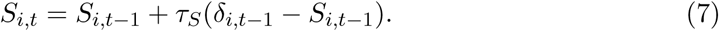

Here, *δ*_*i,t*_ = 1 when an agent chooses option *i* at time *t*; *δ*_*i,t*_ = 0 otherwise. The outcome *R*_*t*_ is equal to 1 when a reward is observed and is 0 otherwise, at time *t*.

### Double Trace model outperforms other RL models in model selection and predictive accuracy

To evaluate a variety of different RL models against the DT model, we performed a model comparison and evaluated the comparative predictive accuracies of the various models. Here, we briefly explain how the DT model compares to the other RL models when computing choice probabilities. These models differ in how they update the value of unchosen options and whether choice history is incorporated into the value update process. The indirect actor model updates the values *Q*_*it*_ for each option *i* in trial *t* for the chosen options only while freezing the values for the unchosen options. The direct model and the *F-Q* model, along with updating the chosen option value, also update the unchosen option value. Thus far, none of these models incorporates choice history as an independent variable. The *F-Q down* model is a version of the F-Q model that discounts the value of the unchosen option with a unique learning rate. F-Q down model was tested with one and two (data not shown) learning rates. F-Q down model with two learning rates is equivalent to the Linear-Nonlinear-Poisson model dveloped by Corrado and colleagues (Corrado et al., 2005) shown by Katahira (Katahira, 2015). We tested the alternative possibility that the F-Q model upcounts (the opposite of discount) the action value of the unchosen option (i.e., the *F-Q up* model). Finally, in addition to reward history, the F-Q with a choice trace (*F-Q W/C*) model also incorporates choice history in the selection of upcoming choices. The F-Q model, with and without a single choice trace, is equivalent to the DT model when *ϑ* = 0 and *ϑ* = *φ* = 0, respectively (see Appendix “RL models” for more detailed descriptions).

First, we tested the performance of different RL models against synthetic data generated by the optimalagents. The DT model outperformed all other tested models (including the F-Q down model with two learning rates) in terms of its predictive power. This was computed as an area under the receiver operated characteristic (ROC) curve, or AUC, for predicted and observed choice distributions. In addition to the AUC, we also used model selection criteria to select the best model (Fig. 3a,b). The model selection criteria were determined based on a cross-validated negative log-likelihood. Briefly, the data were split into five parts, and model parameters were computed on 4/5 of the data. The different models were tested on the remaining 1/5 of the data by computing the negative log-likelihood of the choices (see Methods section on the optimization of parameters for behavioral prediction). We also demonstrate that the DT model was effective at recovering parameters from synthetically generated data by the same DT model with known parameter values. For this, we first generated the data using the specific parameters of the DT model and later using the same model, then extracted parameters from the observed data, as shown in Table 1. Next, we tested various RL models against the mice and human behavioral data. The DT model outperformed all other tested models (Fig. 3a,b). This finding was also evident when looking at the representative choice dynamics that the DT model, mice, and humans generated (Fig. 3c,d).

**Figure 3:**
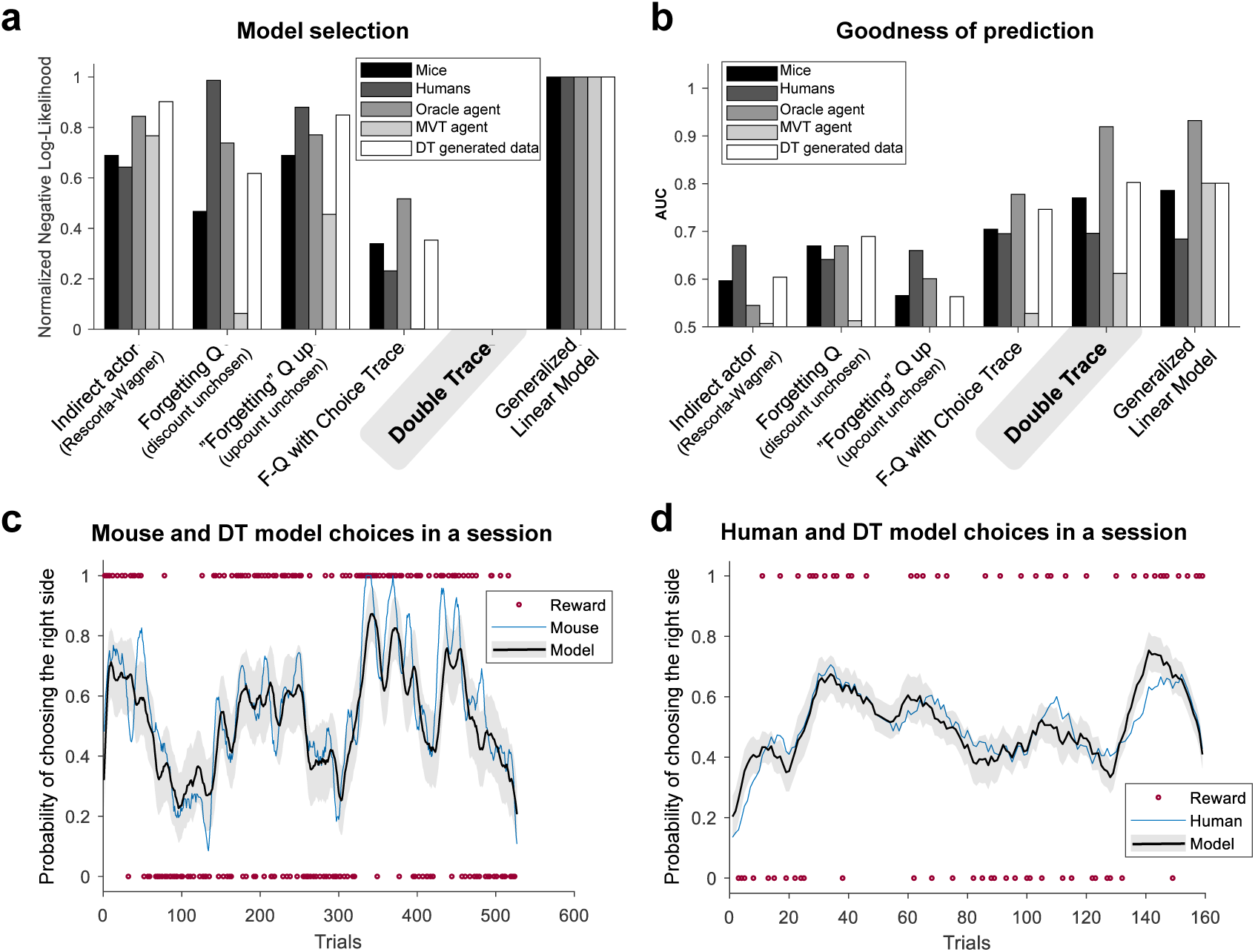
Model selection and goodness of the prediction of mouse, human, and synthetic agent choices. (a) Normalized log-likelihood on test data among models for model selection in predicting the behavioral data of mice, humans, Oracle agent, MVT agents, and DT agents. The direct actor model results were omitted from the plots because their negative log-likelihood tended to infinity. (b) The area under the receiver operating characteristics curve (AUC) on the test data for the goodness of the prediction of the behavioral data of mice, humans, optimal agents, and DT agents. The fitted parameters for mouse behavior using the DT model were *α* = 0.75, *β* = 1.60, *τ*_*F*_ = 0.71, *τ*_*S*_ = 0.24, *ϑ* = 3.31, and *φ* = −2.22. The parameters for human behavior were *α* = 0.84, *β* = 1.77, *τ*_*F*_ = 0.71, *τ*_*S*_ = 0.44, *ϑ* = 1.53, and *φ* = −1.48. With respect to the fit of the Oracle agent behavior, the parameters for the DT model were *α* = 0.19, *β* = 1.60, *τ*_*F*_ = 0.61, *τ*_*S*_ = 0.29, *ϑ* = 10.22, and *φ* = −8.03. The fitted parameters for the MVT agents using the DT model were *α* = 0.89, *β* = 0.23, *τ*_*F*_ = 0.93, *τ*_*S*_ = 0.19, *ϑ* = 23.48, and *φ* = −6.60. For the original and fitted parameters of the DT agents, as well as the boundaries used for the fitting parameters, please refer to Table 1. (c) Example session of the choices made by a mouse and the choices predicted by the DT model; the mean is depicted as bold lines and the standard deviation as shadows. Rewards are depicted as red circles with the value of 1 when the right port was rewarded and 0 when the left port was rewarded. There are no red circles when a trial was not rewarded. (d) Example session of the choices made by a human subject and the choices predicted by the DT model. Rewards are depicted as red circles on 1 when the right port was rewarded and on 0 when the left port was rewarded. There are no red circles when the trial was not rewarded..

Cross-validation tests for model comparison may favor more complex models(Gronau & Wagenmakers, 2019), which may have biased our model selection in favor of the DT model. To avoid this problem, we also performed variational Bayesian model selection (Stephan, Penny, Daunizeau, Moran, & Friston, 2009) using the Matlab toolbox to compute protected exceedance probabilities (https://www.github.com/MBB-team/VBA-toolbox). This analysis revealed that the DT model was the best in all model comparison tests because the protected exceedance probabilities for the DT model were 0.99, 0.75, 0.98, 0.99, and 0.99 when testing against mouse, human, Oracle agent, MVT agent, and DT model generated data, respectively (Suppl. Fig.5).

### The DT model achieves a higher reward harvesting efficiency compared to other RL models

The similarity of choice history effects across species and of the optimal models suggests that the conserved choice history effects are the outcome of the optimization process employed to maximize rewards. If so, then the animal and human characteristic choice history effects captured by the DT model should enable the model to harvest more rewards compared to other RL models. To test the harvesting efficiency of each RL model, we computed regret as a metric. We define regret as the difference between the total rewards *R*_*t*_ collected by an agent and the maximum expected rewards *µ*^*^ (Vermorel & Mohri, 2005). The agent takes action *a* during *T* trials in a session. Thus, regret can be thought of as a simple cost function for evaluating the optimality of a given strategy.

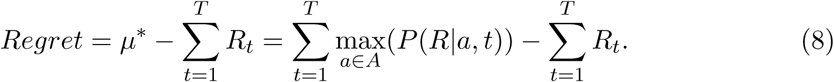

The optimization of the DT model was slightly different than that of the other RL models. Due to the symmetric nature of the two choice traces, whereby different signed parameters can yield the same predictions, we restricted *φ* to negative and *ϑ* to positive values. The constrained DT model performed better than the unconstrained model (data not shown). Therefore, for all of our analyses, we used the parameters of the constrained DT model.

We used the same set reward probabilities in a simulated environment as those used in the mouse experiments (0.1 and. 0.4) and tested the optimized DT model parameters under different volatilities. Our results show that an optimized DT model achieves the minimum regret (Fig. 4a) among all of the tested RL models. It should also be noted that the significant difference between the DT and the next best model occurs at the block length of 100 trials. This block length is close to those used in the animal experiments (Fig. 2a). Furthermore, the slope of the optimized DT models’ reward trace shows a decrement as the block length (inverse of the volatility) increases (Fig. 4b, left panel). This is consistent with previous results (Fig. 2a). Similar changes were observed in the choice trace (Fig. 4b, right panel) with respect to the choice history effects observed in mice (Fig. 2a).

**Figure 4:**
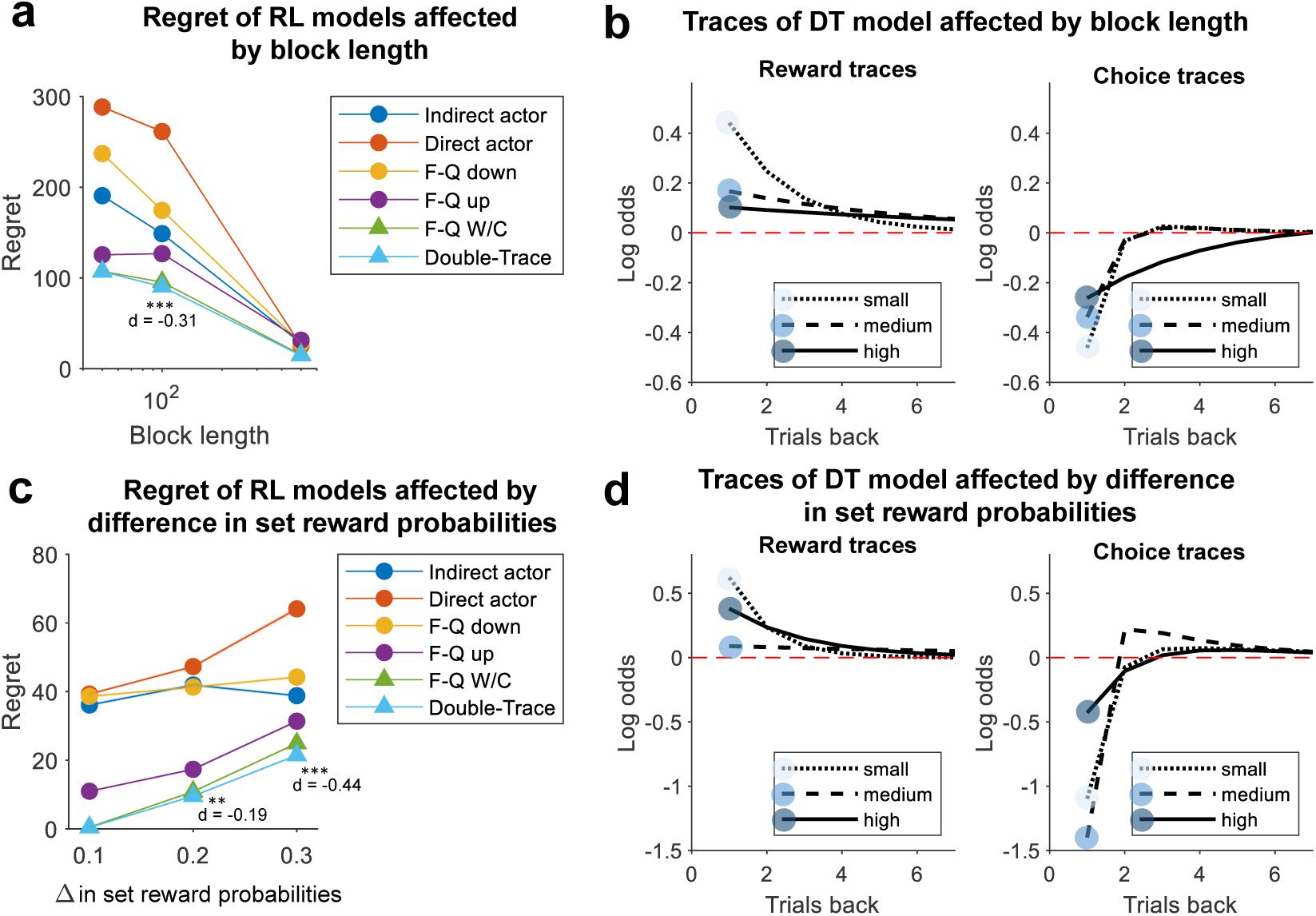
Reward collection by different RL models in the VI task and choice and reward traces of the optimized DT model. (a) The regret for different RL virtual agents as a function of the volatility (block lengths of 50, 100, and 500). The set reward probabilities of the task were 0.1 vs. 0.4. (b) The reward and choice traces of the optimized DT model under the different volatility conditions as described in (a). (c) The regret for different RL models as a function of the difference between the set reward probabilities. The set reward probabilities of the task were 0.45 vs. 0.05, 0.4 vs. 0.1, and 0.35 vs. 0.15. (d) The reward and choice traces of the optimized DT model for the difference in the set reward probabilities as described in (c). The block length was fixed to 100 trials. The regret is defined as the number of rewards away from the optimal collection by the ideal observer in a virtual session with 1,000 trials. The mean is indicated with a solid line, and the standard error of the mean (s.e.m.) is shaded. Note that the s.e.m. is too small to be seen in many cases. Stars represent the p-value of the difference in population medians between the F-Q W/C and DT models using a non-parametric Mann-Whitney U test, in which the DT model showed less regret than the F-Q W/C model (*p* < 0.05 = *, *p* < 0.01 = **, and *p* < 0.001 = * * *). The *d* is the Cohen’s effect size.

When we manipulated the difference in the set reward probabilities as we did for the mice (Fig. 2b), while maintaining the same overall reward probabilities, we found that the DT model outperformed all other tested RL models (Fig. 4c). We also noticed that the choice trace of the optimized DT model increased the short-term alternation component as the difference in the set reward probabilities decreased (0.2 vs. 0.3) (Fig. 4d, right panel). In addition, the shape of the choice trace resembled the choice history effects reported in animals and humans (Fig. 1d-f, Fig. 2b). Contrary to this we failed to see consistent changes in reward trace as a function of difference in set reward probabilities (Fig. 4d, left panel). Based on these results, we conclude that similar to what we observed in mice, the optimized DT model choice trace reflects the growth rates of the rewards on unchosen options.

## Methods

### Rodent behavioural tasks

All the experiments were approved by the Danish Animal Experiments Inspectorate under the Ministry of Justice (Permit 2017−15−0201−01357) and were conducted according to institutional and national guidelines. Water-deprived mice (C57Bl/6J strain) were trained to initiate the trials by poking their noses into a central port equipped with sensors that detect the exact entry and exit of animals (Fig. 1 b). After poking their noses into the central port, the mice were free to poke their noses into the left or right sides of the port. Rewards were delivered according to the VI schedule following eq.1. There was no fixed inter-trial interval. Mice could initiate the next trial immediately after poking their noses into one of the sides of the port. We did not use any punishment or time-out period. 267 sessions using eleven mice (18 − 47 sessions per mouse) were recorded.

### Human behavioural tasks

The VI task in the form of a computer game was performed by 19 individuals (aged 18-60). The virtual task environment consisted of two apple trees and a gate separating the avatar from the trees (Fig. 1 a). The player controlling the avatar had to first open the gate (delay intervals: 0-1s, 2-3s and 4-5s) before subsequently choosing one of the trees. On each trial apple tree provided one apple with the probability that followed the VI task schedule(eq.1). Each participant received brief instructions (see Appendix) on how to play the game. They were, however, not informed about the specifics of the reward schedules or the task structure. Each participant completed at least six blocks consisting of 20-30 trials. Within each block, the task conditions (gate intervals and reward probability) remained constant but changed randomly between blocks.

### Optimal agents

While optimal agents do not necessarily offer biologically realistic models of choice, they are still informative both conceptually, and it terms of providing upper bounds of performance given their specific constraints and assumptions. We constructed the following three optimal agents:

### The Oracle agent

The generation of choices of a optimal agent follow a simple rule: Choose the option *i* with the highest current probability of receiving a reward, when *i* = *argmax*_*a*∈*A*_*P*(*R*|*a, t*). The update of this probability follows eq. 1 when there are no block transitions. However, if set reward probabilities change, during block transitions, then the rule is:

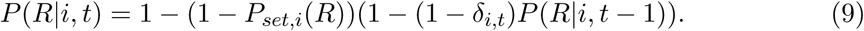

*δ*_*i,t*_ = 1 when an agent chooses option *i* at time *t*; *δ*_*i,t*_ = 0 otherwise. This agent runs in a virtual session of *T* = 1000 trials.

### The Bayesian agent

The Bayesian agent was constructed to infer the set reward probabilities *P*_*set*_(*R*|*i, t*) of each side *i* ∈ {1, 2} at trial *t* = 1, 2, …, *T* while taking actions and observing rewards *R* ∈ {1, 0} in a VI task. We assume that the agent knows the structure of the baiting equation given by the VI environment (eq. 1) and follows that equation to update their estimates. We ran this agent by initializing with *N* = 1000 random estimates of the set reward probabilities 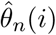, where *n* = 1, 2, …, *N*, drawn from a uniform probability distribution 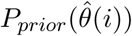 with boundaries at 0 and 1. This agent ran in a virtual session of *T* = 1000 trials or iterations, where at every trial *t*, it estimated the reward rates,

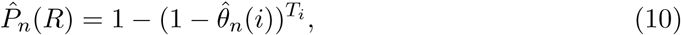

where *T*_*i*_ is the number of trials since the option *i* was chosen. Then, it generated the value of each option as

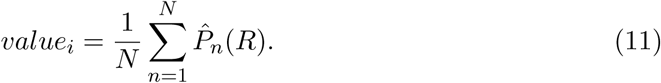

The Bayesian agent followed a greedy policy, meaning that it choose the action *i* with the highest value at *t* with respect to the other action (∼ *i*); in other words,

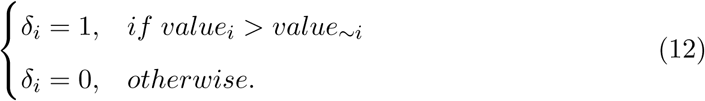

After an action is taken, a reward *R* = 1 or a no reward *R* = 0 is observed by the Bayesian agent. The likelihood of updating the priors will then be

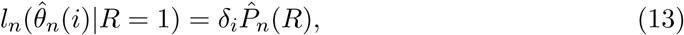

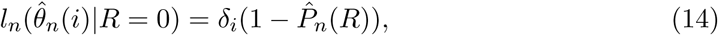

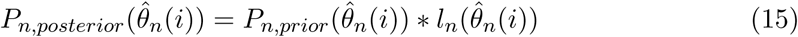

The multiplication of the prior probability of estimated set reward probabilities 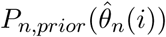 with the likelihood will lead to the posterior probability of estimated set reward probabilities 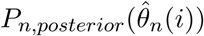. The sum of the posterior probabilities is then normalised such that it sums to one. The new estimated set reward probabilities are sampled randomly from the range of posterior probabilities to update the old estimates. The posterior probabilities become then the prior probabilities for the next trial. A noise factor of +*/*−2.5% or *ϵ*_*n*_, drawn from a uniform probability distribution with boundaries [-0.025,0.025], is added to the new estimates of the set reward probabilities 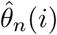. This noise factor helps the model to explore wider range of possible set reward probabilities and converge more precisely to the true set probabilities.. Since the noise term *ϵ*_*n*_ is added to the new estimates of the set reward probabilities, it is possible that some of these new estimates 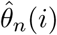 fall outside of the original boundaries of 0 and 1. We corrected for this as follows:

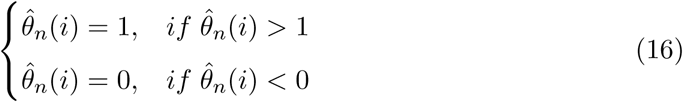

A new iteration starts with trial *t* = *t* + 1 until the session is over at the end of the last trial *t* = *T*.

### The MVT agent

An environment with heterogeneous patches was created following work by Bettinger L. and Grote N. (Bettinger & Grote, 2016). The maximum amount of reward per patch *G* was drawn from a uniform distribution *G* ∈ [0, *G*_*max*_]. A reward decay rate for each patch was drawn randomly from an exponential distribution with a mean of 0.25. The handling time ∈ [0 2] and the travel time-*tt*_*i*_ ∈ [50*s ttravel*_*max*_] were drawn also randomly from a uniform distribution.

It is important to mention that to follow the algorithm proposed in (Bettinger & Grote, 2016), it is necessary to compute the average net intake rate. This was done by considering a heterogenous environment as in (Calcagno, Mailleret, Wajnberg, & Grognard, 2013), and the average net intake rate was computed by

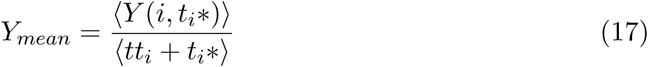

as the mean of reward rates *Y* (*i, t*_*i*_*) given by each *i* patch with its optimal stay time *t*_*i*_*. This optimal stay time in a patch is found when the marginal reward value in a patch <u>^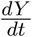^</u> matches with the reward obtained in that patch *Y* (*i, t*_*i*_*) over the total time spent on that patch, i.e., the addition of the stay time *t*_*i*_* and the travel time *tt*_*i*_, as originally proposed by Charnov (Charnov, 1976).

### Linear regression for computing reward and choice history effects

The influence of the past rewards *R*(*t*−*n*) and choices *C*(*t*−*n*), with *n* = 1, 2, …, 15 trials, on the upcoming choice *C*(*t*) of an agent was represented by the coefficients 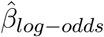 in the linear domain of a logistic regression, also known as log odds. To calculate the coefficients, we defined the reward vector as the difference in experienced rewards between options as *R*(*t*) = *R*_*Right*_(*t*) − *R*_*Left*_(*t*) at every trial *t*. The rewards take values *R*_*Right*_(*t*) = 1 when the reward was delivered and that trial or *R*_*Right*_(*t*) = 0 when subjects choose right option and did not recieved a reward or if left option was chosen. The inverse was true for *R*_*Left*_(*t*).

The choice vector was defined as *C*(*t*) = 1 if the right option was chosen at trial *t* and *C*(*t*) = 0 otherwise. The logistic regression was then regularised with an elastic net that linearly combines penalties *L*^1^ and *L*^2^ from lasso and ridge regression (Hastie et al., 2009). The penalty term of the elastic net is defined as *λP*(*β*_*log*−*odds*_), where *P* is the number of parameters (*β*_*log*−*odds*_) and interpolates between the *L*^1^ and *L*^2^ norms as follows:

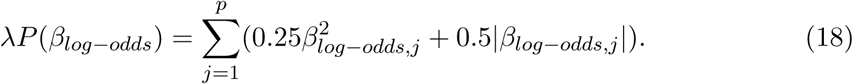

Therefore, the regression coefficients that had the minimum mean deviance (or, equivalently, the maximum mean log-likelihood) as a function of the tuning parameter *λ* in a five-fold cross-validation process were selected as 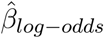 We imlemented logistic regression in MATLAB (www.mathworks.com) using *lassoglm* function.

### Optimisation of parameters for behavioural prediction

The choice history *δ*_*i*_(1, 2, …, *t* − 1) and reward history *R*(1, 2, …, *t* − 1), were introduced to various RL models as *x*_*t*−1_ to compute the action values and determine the probability of choosing an action in the current trial *t* using the softmax action selection rule:

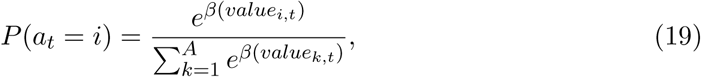

where *value*_*i,t*_ is determined by the model used (more details in the Appendix).

This probability *P*(*a*_*t*_ = *i*) and the actual action *δ*_*i,t*_ at time *t* with the set of parameters *θ*_*RL*_(*n*) per model determine the log-likelihood as follows:

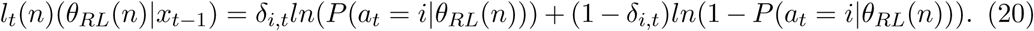

To find the optimal parameters 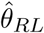 for each model, we initially selected 1,000 combinations *θ*_*RL*,1_ from a uniform distribution, where the boundaries of the parameter space were *α* ∈ [0, 1], *β* ∈ [0, 50], *φ* ∈ [−25, 25], *ϑ* ∈ [−25, 25], *ϑ* ∈ [−25, 25], *τ*_*F*_ ∈ [0, 1], *τ*_*M*_ ∈ [0, 1] and *τ*_*S*_ ∈ [0, 1]. Next, we took 1% of the combinations with the maximum mean log-likelihood and used the mean 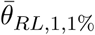 and standard deviation *σ*(*θ* _*RL*,1,1%_) of this subset to draw a new set of 1,000 combinations as follows:

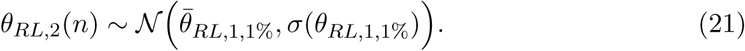

We repeated this process several times to narrow the original parameter space until the highest log-likelihood (a negative value or zero) was less than 99.99% of its value for two iterations in a row. The optimal parameters for the prediction of each model 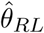 were then the combination of parameters with the highest mean log-likelihood of the last iteration in the optimisation process.

The optimised parameters for each model and the estimated coefficients were trained and tested via five-fold cross-validation on the behavioural data in order to obtain the average of the minimum negative log-likelihood and the average AUC. These latter metrics were used for model selection and the goodness of prediction for each model, respectively. To compute the AUC, the probability 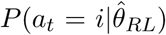 was set as the score and the original action *δ*_*i,t*_ as the label. We used Matlab function *perfcurve* to compute the AUC scores.

### Optimisation of parameters in simulated environments

Regret (eq. 8) was used to estimate the optimal parameters for each model in the VI task We iteratively determined the best parameters for each model by selecting a random set of initial parameters *θ*_1_ uniformly 10,000 times and by simulating one session of *T* = 1000 trials four times. We tested two conditions, the first one is the VI task under different degrees of volatility (i.e. block sizes): *N*_*bl*_ ∈ {50, 100, 500, 1000} and one pair of the set probabilities 0.10:0.40. The second condition of the VI task have three different pairs of set probabilities 0.20:0.80, 0.30:0.70 and 0.40:0.60, and one block size *N*_*bl*_ = 100. The boundaries of the parameter space were *α* ∈ [0, 1], *β* ∈ [0, 10], *φ* ∈ [−2, 0], *ϑ* ∈ [0, 2], *τ*_*F*_ ∈ [0, 1] and *τ*_*S*_ ∈ [0, 1]. Since the number of combinations was insufficient for a thorough search of the parameter space, we selected the 5% with the lowest regret 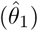 out of the original 1,000 combinations.

We thereby realised a local search algorithm in our multidimensional parameter space, extending three standard deviations around the top 5% of the previous parameter combinations and selecting a new random set of parameters *θ*_2_ 5000 times:

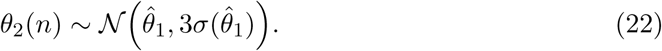

We repeated the selection of best parameters by for *θ*_2_ the same way described for *θ*_1_. It is important to mention that at every iteration, a different random number generator was selected to prevent running the generator under the same probabilities during each trial and to give the model the opportunity to explore different probabilistic scenarios. However, this does not guarantee that the combinations had low regret due to this random selection. To overcome this potential problem, we took the 10% of *θ*_2_ with the lowest regret, having 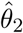, and tested each combination 100 times under different scenarios to determine consistency of the same outcomes. The parameter combination in 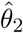 with the lowest regret on average was selected as optimal.

## Discussion

Optimal models of animal behavior formulate rules that an agent should follow to maximize some currency that ultimately determines or contributes to fitness (Charnov, 1976; McNamara & Houston, 1986; Stephens & Krebs, 1986). When foraging, it is typically assumed that this currency is the net reward rate. However, evaluating whether animals follow the optimal foraging principles faces several challenges. This is because in experimental settings that approximate natural foraging, many variables such as energetic, time, and opportunity costs are difficult to measure, and competing models can generate qualitatively similar predictions (Walton, Rudebeck, Bannerman, & Rushworth, 2007; Hayden, Pearson, & Platt, 2011; Geana & Niv, 2014). Therefore, it is not known whether such optimality models are adequate for explaining animal behavior, and if so, how these models can be generalized to ecologically relevant habitats. The matter is even more complex for perceptual decision tasks, because the optimal inference of perceptual objects is complicated by the neural hardware operating under stringent biological constraints (Beck, Ma, Pitkow, Latham, & Pouget, 2012). Therefore, suboptimal inference is compatible with both behavioral variability and suboptimal performance in perceptual decision-making tasks (Tjan, Braje, Legge, & Kersten, 1995). These problems can be mitigated in reward foraging tasks that minimize the complexity of the task-relevant perceptual objects and instead require subjects to detect either the presence or absence of past rewards to guide future choices. Furthermore, in a large class of two alternative choice tasks, the test animals initiate decisions from identical starting points to equally distant options. Therefore, it is fair to assume that different choices typically embody the same costs on a decision maker. Because of the property of two alternative choice tasks in this study, we were permitted to formulate optimal models of behavior.

Viewed from the optimization perspective, we now explore the foraging decisions of animals and humans. Namely, we address the contingencies between behavior and reward availability, and how these may configure choice history effects. We examine this question from the narrow perspective of biological and synthetic agents that attempt to maximize their reward rates. First, we show empirically that choice history effects track the changes in reward probabilities for unchosen options in a multiple choice task. This effect is distinct from reward history effects, which are predominantly associated with the volatility of the reward probabilities. Choice history effects are also distinct from the reaction times, which (Behrens et al., 2007; Genest, Stauffer, & Schultz, 2016; Kheifets, Freestone, & Gallistel, 2017) reflects the overall reward rates averaged over the options. By showing the same choice history effects in optimal models and in three different species, we argue that these choice history phenomena are generally adaptive in habitats where past consumption of rewards effects the availability of rewards in the future. We incorporate choice history into a new RL variant (Sutton & Barto, 1998) that, when compared to existing models, had a higher predictive adequacy for biological behavior and in synthetic agents yielded higher reward rates, and thus energy harvesting efficiency. Finally, we generalize this finding to show that the choice history effects observed in two alternative choice tasks are evident in optimal agents that forage in habitats with diminishing energetic returns.

Here, we note a number of limitations that accompany our studies and offer counter-arguments that may mitigate these concerns. Though arguably more ecologically valid than experiments in which choice history is decoupled from reward availability, our task could still be limited in its ecological validity. This issue could be remedied by employing more naturalistic foraging fields and by affording greater degrees of motoric freedom when foraging. A number of studies in rodents and humans have explored more ecologically realistic scenarios with innovative task designs (Hayden et al., 2011; Walton, Bannerman, & Rushworth, 2002; Wikenheiser, Stephens, & Redish, 2013). However, the benefit of the more artificial experimental setup of the VI task is the precise control of reward probabilities, while also controlling energetic costs and motoric uncertainties. Estimating the cost of travel or time in foraging animals is difficult, which makes it challenging to compare the performance of the animals with that of the optimal agent models. Here, we chose to focus on a simple and straightforward behavioral task design and used the model derivation to generalize to ecologically relevant contexts.

### The behavioral function of reward and choice history effects

Animal behavior must be tuned to environmental statistics to maintain its adaptive function(Genest et al., 2016; Kheifets et al., 2017; Behrens et al., 2007). Different dynamics can be discerned in natural habitats that are expected to shape foraging decisions. For example, one such dynamic is that the forager experiences volatility in its food reserves or in the availability of food in its habitat. Another is that there is either positive of negative growth of food availability at unvisited patches. Both of these processes should be incorporated into the decision process to generate adaptive behavior. The regression-based models as well as RL-based models that incorporate experienced reward statistics or reward history into the decision process (Corrado et al., 2005; Sugrue, Corrado, & Newsome, 2004; Bari et al., 2019; Kim et al., 2009) adequately describe both an animal’s choices and neural responses. However, these models are well suited to deal with volatility (Behrens et al., 2007) when unchosen options remain static. Indeed, in VI type habitats, these models (Katahira, 2015) (F-Q is one of those models) can be substantially improved when choice history is also incorporated into the decision process. Thus, we argue that reward and choice history effects reflect complementary decision processes tuned to capture the wider dynamics of the natural habitats.

If choice history effects emerge as an evolutionary adaptation to natural habitats, then the full suppression of this behavior, even in well-trained animals, in a perceptual or value-based decision-making task might be difficult (Akrami et al., 2018; Hwang et al., 2017). Indeed, in contrast to perceptual decision-making tasks that expose animals to a static environment, models that assume a dynamic environment are better at capturing the animals’ choices (Mendonca et al., 2020). However, when an environment is not static and task-relevant stimuli exhibit autocorrelations, choice history effects carry an important adaptive function (Braun, Urai, & Donner, 2018). The experimental data showing that neural representations of choice history are widely distributed in sensory, motor, and association areas (John-Saaltink, Kok, Lau, & De Lange, 2016; Pape & Siegel, 2016; Hwang et al., 2019) suggests that these representations carry adaptive function.

### Implications for neural representations

The existence of short-term alternation and long-term perseverance in choice history effects that can be described by two time-scale exponentials raises other questions regarding neural representations. Are slow and fast processes encoded by distinct neural populations? Or do the same neurons represent two variables that emerge over different time-scales? The two time scales of decision processes appear in other reward foraging and perceptual tasks. In sequential decision-making tasks, normative models based on Bayesian principles (Bogacz, Brown, Moehlis, Holmes, & Cohen, 2006) reveal within- and across-trial accumulation processes that operate on two different time scales (Glaze, Kable, & Gold, 2015). The two time scales of value accumulation also appear in optimal models with foraging tasks that provide fixed (i.e., non-replenishing) reward probabilities (Iigaya et al., 2019), albeit with different timescales. Thus, fundamental questions emerge as to how individual neurons capture the multiplexed nature of decision-making phenomena.

### Benefits of choice history effects for a wider class of foraging tasks

What benefits could choice history effects and its implementation in the form of an RL model provide to foraging agents that navigate natural habitats that may not be limited to only two options? MVT specifies rules for optimal foraging in habitats with an arbitrary number of food patches (options) that provide diminishing returns due to finite resources (Charnov, 1976). MVT provides abstract decision rules for optimal agents in a steady state, where the agent knows the average reward rate over the entire habitat. In contrast to agents that adhere to MVT, RL agents can, in principle, learn the average rate of rewards of the habitat by exploration and experience. However, studies that have compared MVT agents and RL performance(Constantino & Daw, 2015) showed that the agent that follows the temporal difference (TD) model harvests less energy than the agent that follows the MVT rule in habitats with monotonically diminishing returns. Thus, the advantage that the MVT rule provides over the TD model comes at the expense of the flexibility to adapt to environmental contingencies that deviate from the habitat to which the MVT agent is adapted. In contrast to the TD models, the DT model is well positioned to forage in habitats with diminishing returns by deploying choice history effects into the decision process. This fact enables DT agents to forage efficiently in more ecologically realistic habitats.

## Conclusion

Our results show that choice history effects in species as diverse as mice, monkeys, and humans can be phylogenetically conserved, which lends credence to the hypothesis of a shared evolutionary-selected mechanism. The perspective offered here suggests that the choice history effects evolved as a behavioral adaptation to habitats where one’s past choices can contribute to diminishing returns. We provide an explanatory and algorithmic account of choice history effects beyond their description as a bias that connects this concept to a broader class of optimality models within behavioral ecology.

## Appendix

### Derivation of the DT model from logistic regression

#### Logistic regression model of behaviour

The probability that a foraging animal will choose the left side in a discrete version of a VI task with two choices is *P*(*a*_*t*_ = *l*). Lau and Glimcher (Lau & Glimcher, 2005) originally proposed to compute the probability *P*(*a*_*t*_ = *l*) as a linear combination *h*_*l,t*_ of the reward *R* and choice *C* history of the last *M* trials with their corresponding coefficients *b*_*R*_, *b*_*C*_, plus a constant bias *K*. As the choices are binary in nature, the linear function *h*(*t*) is then transformed in logit space to a probability, as shown in Equation 20. Here, *β* is the exploration factor. This is also known as a log-linear model or logistic regression:

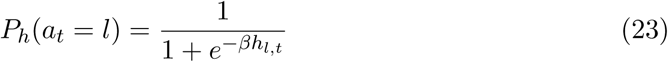

and

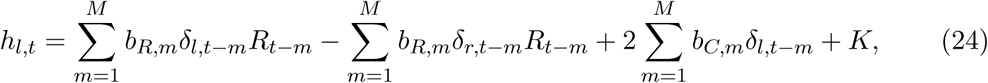

where the chosen action *i* at time *t* is depicted as *δ*_*i,t*_ = 1.and the reward at time *t* is described as *R*_*t*_ = 1. *K* is the bias term in the equation.

#### Decay in the reward and choice history regression coefficients

The results of our experiments and previous studies (Lau & Glimcher, 2005; Rutledge et al., 2009; Bari et al., 2019) *b*_*R,m*_ and *b*_*C,m*_ decay, as a function of the *m* trial history (Fig. 1 g-i). We propose that these decays for the reward and choice history to be modelled according to the following equation:

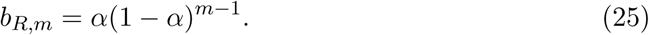

Here, *α* characterises the decay rate for rewards. *τ*_*F*_ and *τ*_*S*_ characterise the decay rate for choices according to:

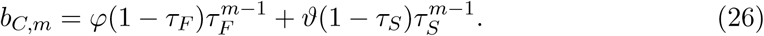

*φ* is the scaling factor for choice trace *F*, and *ϑ* is the scaling factor for choice trace *S*.

The function *h*_*t*_ of the logistic regression was separated into reward and choice terms to ensure simplicity in the next analysis.

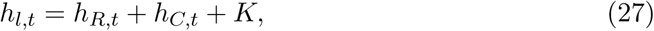

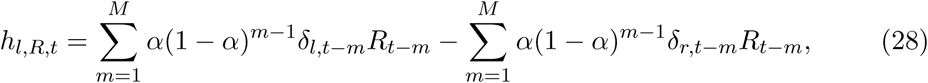

and

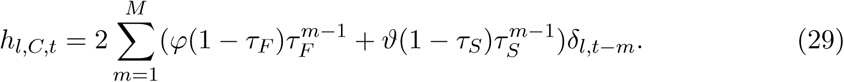

In the next section, we show how these equations were used to derive the recursive DT update rule:

#### From logistic regression to the softmax selection rule

Like the logistic regression model (eq. 23 and eq.24), the RL model computes choice probabilities using the value expectation of rewards (Q, eq. 4) under the softmax selection rule as follows:

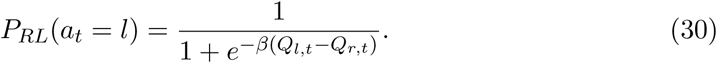

*β* defines the inverse temperature (also called the exploration factor), and *Q*_*i,t*_ is the reward expectation value of a given action (or option) *i* at time *t*. In this case, *l* and *r* refer to the left and right options, respectively.

We propose that one way to reconstruct the animals’ choice history effects (Fig. 1 g-i) within the RL framework is to incorporate a separate double-choice history component in the softmax selection rule. Using the difference between the two choice traces, each computed with a different learning rate, we can recover the characteristic (wavy) shape of the animals’ choice history effects. Here, we use learning rates that are independent, such as *τ*_*F*_ and *τ*_*S*_, and that can generate a vast diversity of curves. The softmax function then takes the following form:

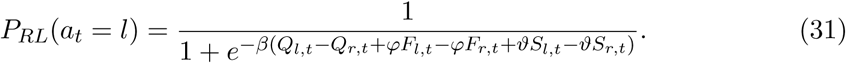

Here, *F* is the quickly decaying choice trace (eq. 6), and *S* is the slowly decaying choice trace (eq. 7).

Next, we show the equivalence of the DT model (eq. 30) with the logistic regression model (eq. 23). Therefore, the following equivalence should be satisfied:

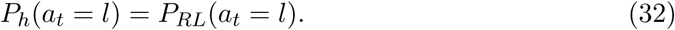

It follows from this equivalence that

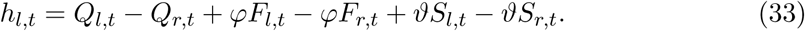

#### Reward expectation and its equivalence in logistic regression

For the reward expectation, it was already shown that the logistic regression model is equivalent to the F-Q model (Katahira, 2015), which is characterised by the following update rule:

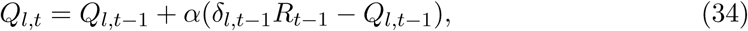

where *α* is the learning rate.

Katahira (Katahira, 2015) proposed that the equivalence in eq. 33 can be reduced to eq. 34:

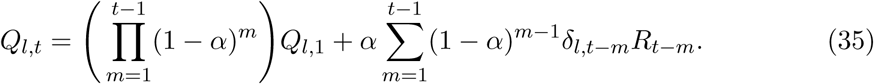

There are enough trials *t* >> 1 to neglect events that occurred in the very distant past (Katahira, 2015):

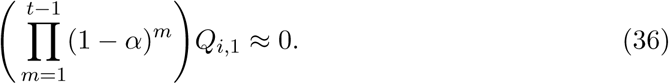

Under this assumption, we define *t* − 1 ≈ *M*, and, therefore, it is possible to observe how this update rule for the reward expectation is equivalent to the reward coefficients in logistic regression:

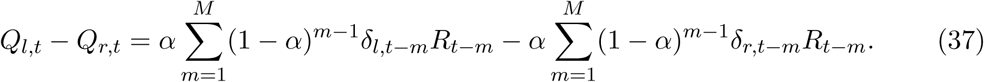

Therefore,

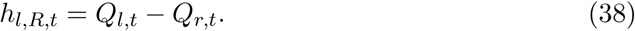

#### Choice traces and their equivalences in logistic regression

In the next section, we demonstrate how to reconstruct the proposed choice history decay observed in the experimental data in a recursive manner:

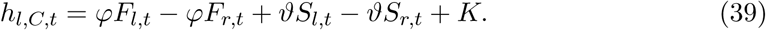

We introduce a recursive function that we call a choice trace, and its equivalence in logistic regression is shown in the next three equations. We follow similar derivations as those of Katahira in his work (Katahira, 2015). We introduce the functions fast (F) and slow (S), corresponding to the fast and slow learning rates in the update rules, respectively (Equations 6 and 7).

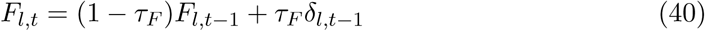

We show the equivalence of the choice trace with the logistic regression in the following derivation:

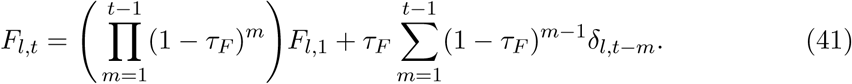

Under the same assumption as that of the reward expectation, when the number of trials is high enough *t* >> 1 that very distant past events become negligible, we can reduce the previous equation as follows:

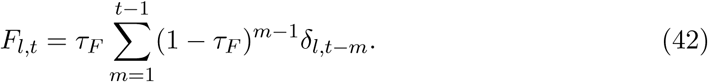

In the next equations, we can observe how the choice trace of the chosen side is the same as that minus the choice trace of the unchosen side in a two-alternative choice task.

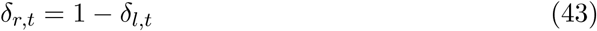

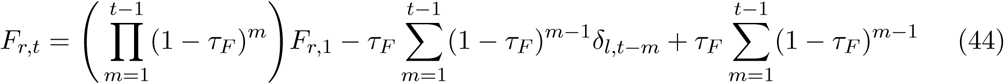

When *t* >> 1,

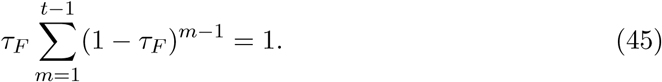

Finally,

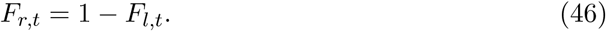

Therefore, the difference between the two choice traces with the same learning rate can be represented as follows:

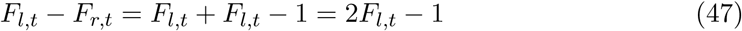

and

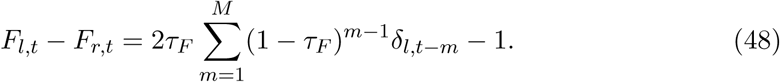

The second choice trace is called slow because the learning rate is slower than that in the previous choice trace by definition. The equivalence of this recursive equation with a sum and a decay follows the same logic as the previous choice trace.

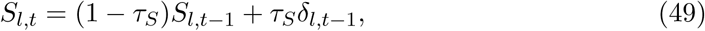

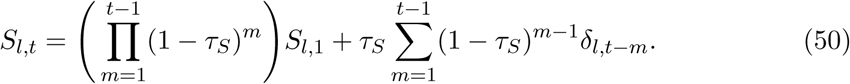

Assuming that *t* >> 1,

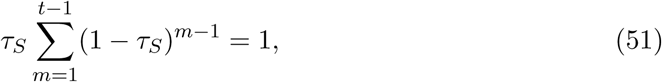

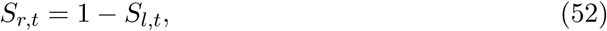

and

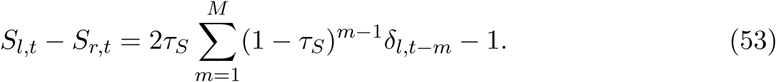

Based on the choice part of Equation 25, we obtain the following equations:

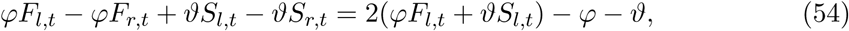

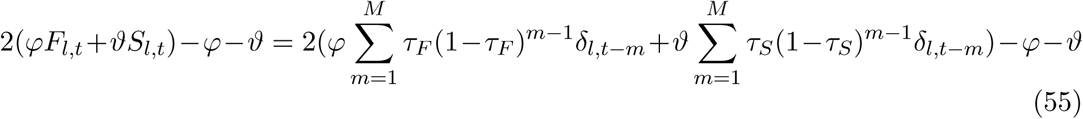

and

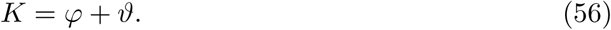

We then show how, using the fast and slow choice traces, we can recover Equation 28 as a part of the original logistic regression, where *φ* and *ϑ* are the scaling factors. These factors allow the agents to select which choice trace, fast or slow, should dominate the decisions.

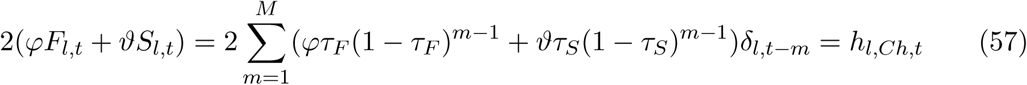

Thus,

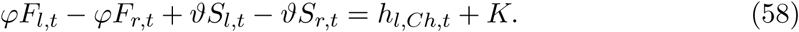

#### RL models

##### Indirect actor model

The indirect actor model updates the reward expectation (or state-action value) Q only for the chosen option.

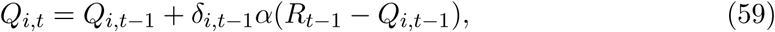

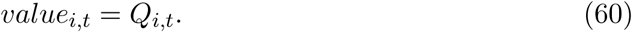

##### Direct actor model

The reward expectation of the direct actor model is updated based on the probability of the chosen action and the reward outcome. This rule also affects the reward expectation value of the unchosen actions.

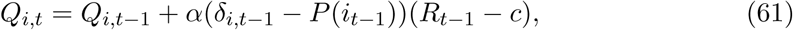

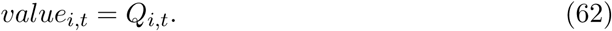

##### F-Q down model

The F-Q model is a slight modification of the indirect actor model, where the reward expectation value of the unchosen actions are forgotten and vanish to zero.

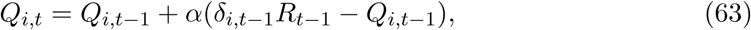

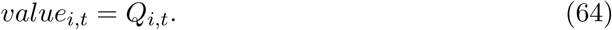

##### F-Q with the choice trace model

According to the F-Q W/C model, the probability of taking the next action will depend not only on the reward expectation but also on choice trace F updated according to the following:

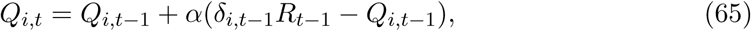

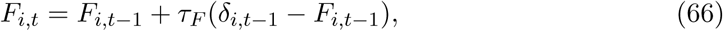

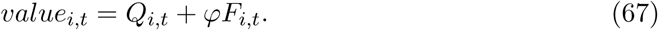

##### F-Q up model

While the F-Q model decreases the value of the unchosen actions, the proposed model increases their value up to *C* with its own learning rate *α*_*up*_, which acts as a positive counter for unchosen actions that adds to the action value.

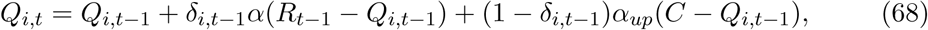

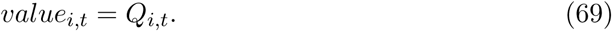

#### The exact instructions given to the subjects who participated in the human-VI task

The following text is the instruction (varbatim) given to the subjects who participated in human-VI task: “In this experiment you will play a game about collecting apples. The game takes around 20-30 min to complete. When playing this game please choose a place with little distractors around for the time of the game. Please play until a notification tells you that you have completed enough rounds. After that you may continue playing or quit the game. To quit, use the “escape” key. After you have finished all rounds, there will be a short questionnaire. If you wish to end the game before you have completed all rounds, use the “escape” key. You will not be directed to the questionnaire and your data will not be used. In the game you play an avatar who collects apples from two identical-looking trees. To begin the game, you need to hold down the “space” key until the gate in front of the two trees opens. After the avatar has stopped moving, use the arrow keys to choose between the trees. A falling apple will visualize whether you received an apple in that round.”

#### Notations and symbols

##### General notation

*t*: Time (trial)

*T* : Number of trials

*i*: Action (option)

*a*_*t*_: Chosen action (option) in trial *t*

*A*: Number of actions (options)

*R*_*t*_: Observed reward at time *t*

*δ*_*i,t*_: Action *i* at time *t* delta function

##### Logistic regression and reinforcement learning

*l*: Left option

*r*: Right option

*m*: Past trial

*M* Number of past trials

*K* Constant term

*h*_*i,t*_: Predictor for action *i* at time *t*

*h*_*i,R,t*_: Predictor for action *i* at time *t* using only the reward history

*h*_*i,C,t*_: Predictor for action *i* at time *t* using only the choice history

*b*_*R,m*_: Reward coefficient for *m* trials back

*b*_*C,m*_: Choice coefficient for *m* trials back

*P*_*h*_(*a*_*t*_ = *i*): Probability of taking action *i* at time *t* using function *h*

*P*_*RL*_(*a*_*t*_ = *i*): Probability of taking action *i* at time *t* using an RL model

*α*: Reward learning rate

*τ*_*F*_ : Choice learning rate (fast)

*τ*_*S*_: Choice learning rate (slow)

*β*: Inverse temperature (exploration factor)

*Q*_*i,t*_: Reward expectation (state-action value) of taking action *i* at time *t*

*F*_*i,t*_: Fast choice trace of action *i* at time *t*

*S*_*i,t*_: Slow choice trace of action *i* at time *t*

*φ*: Scaling factor for the fast choice trace

*ϑ*: Scaling factor for the slow choice trace

*c*: Constant term in the direct actor model

*value*_*i,t*_: Model dependent value of option *i* at time *t*

⟨⟩: Average

*Y* (*i, t*_*i*_): Rewards collected in the patch *i* at time *t*_*i*_

*tt*_*i*_: Travel time to patch *i*

*t*_*i*_: Time foraging in patch *i*

*ttravel*_*max*_: Maximum travel time possible to get to a patch

*G*_*max*_: Maximum pre-pristine factor: Maximum density of rewards in a patch

*G*: Pre-pristine factor: Density of rewards in a patch

##### Optimisation

*µ*^*^: Maximum number of expected rewards for the ideal observer

*θ*_*step*_(*n*): Combination *n* of parameters in a given step of the optimisation method

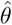: Best parameter combination found after optimisation

*θ*_*RL*_(*n*): Combination *n* of parameters for a given RL model

*l*_*t*_(*n*)(*θ*_*RL*_(*n*)|*x*): Log-likelihood of parameters *θ*_*RL*_(*n*) given *x* data

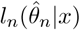: Log-likelihood of parameters 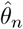 given *x* data

*k*: Number of parameters

*N* : Number of starting parameter combinations

#### Implementation

The analyses of the behavioural and computational models were performed using MAT-LAB (MathWorks, Inc.). The code is available upon request.

## Acknowledegments

We thank Larry F. Abbott and Ashok L. Kumar for their suggestions on the DT model and the manuscript, Naoshige Uchida for his suggestion to highlight the non-Markovian property of the VI task. We thank Sophie Seidenbecher and Madeny Belkhiri for their assistance with editing the manuscript. We thank Søren Rud Keiding for his advice and discussions, Eske Nielsen for programming the human game in Unity platform and Maris Sala and Daniel Kozlovski for assisting with the data collection. This work was supported by Lundbeckfonden, grant no. DANDRITE-R248-2016-2518. Junior Samuel López-Yépez was supported by the Lundbeck PhD Fellowship, grant no. R191-2015-1506.

## Supplemental Figures

**Supplementary Figure 1:**
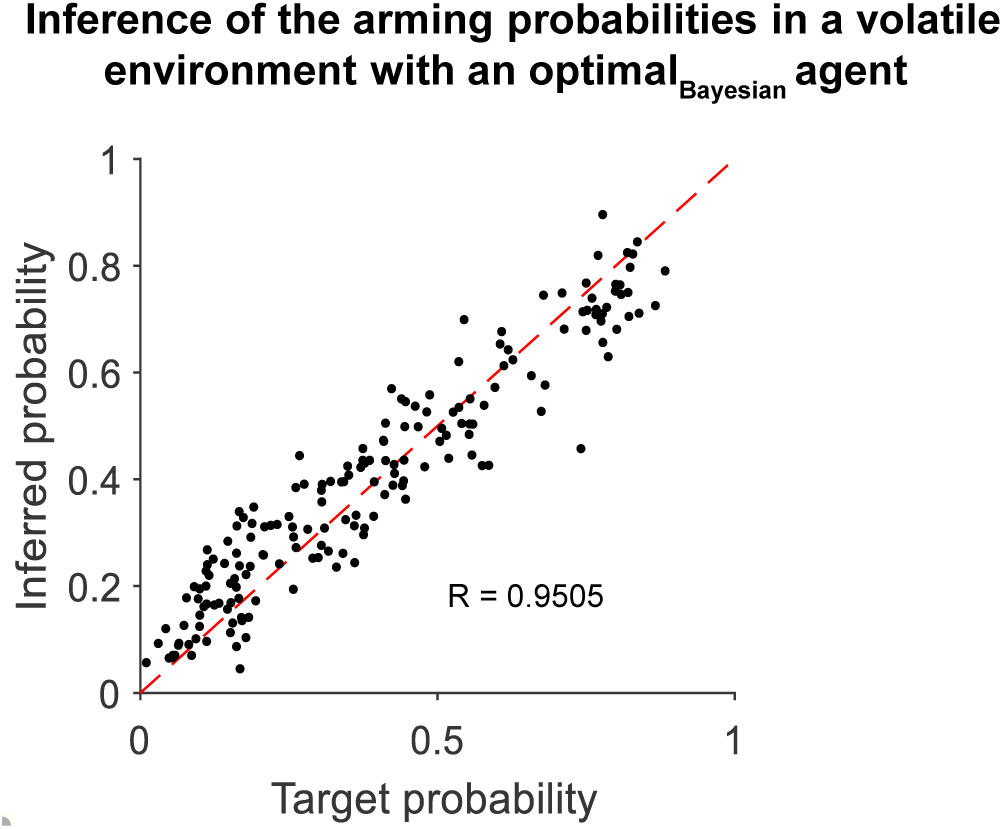
Inference of set probabilities of a Bayesian agent. Correlation of the inferred probability and the set probabilities by a Bayesian agent (see methods) in the last trial of the session. Each point represents a simulation (n = 90) of a session with *T* = 1000 trials, where the block size was kept to 100 trials.

**Supplementary Figure 2:**
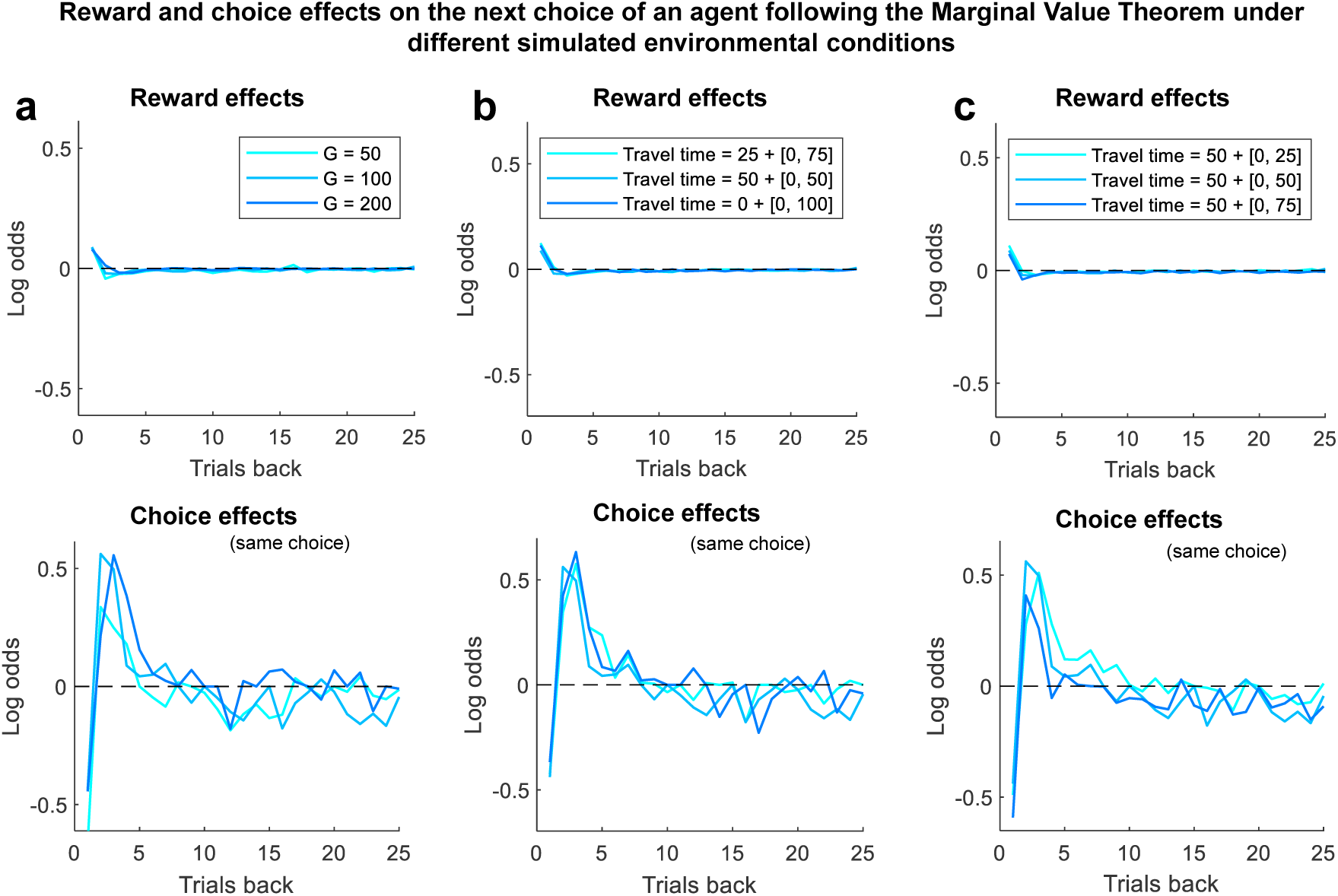
Reward and choice influence in heterogenous patches. The influence of past rewards and choices on the upcoming decision is stable when the MVT is run in simulated environments that vary in the pre-pristine factor G (a), the total travel time (b) and the uncertainty in the travel time (c). The range [x, y] represents a value drawn from a uniform distribution with boundaries x and y. Each condition was run 1000 times, and only the mean of the coefficients (log odds) are shown. The log odds were obtained through logistic regressions with elastic-net regularization with 5-fold cross-validation.

**Supplementary Figure 3:**
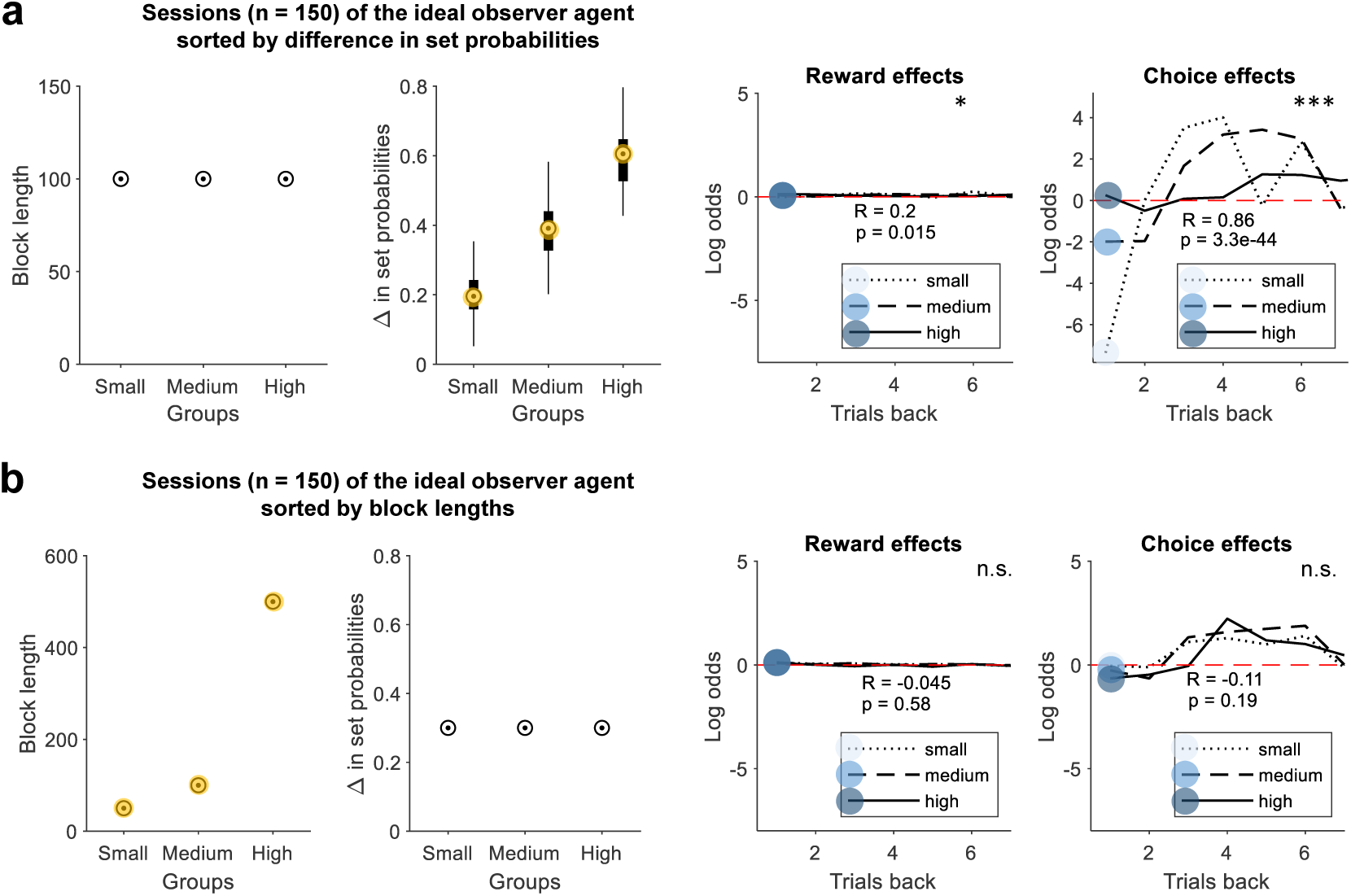
Change in the difference of set reward probabilities affects the choice history effects of the Oracle agent. (a) Sessions of the Oracle agent sorted in three groups by the difference (Δ) in set reward probabilities (150 sessions with 1000 trials each). The set reward probabilities pair per session were 0.4 vs 0.6, 0.3 vs 0.7 and 0.2 vs 0.8 as the means with a standard deviation of 0.064. The two left panels show the distribution of the block lengths and the difference in set reward probabilities. The two right panels show their correspondent reward and choice kernels. (b) Sessions of the Oracle agent sorted in three groups by the difference (Δ) in set reward probabilities (150 sessions with 1000 trials each). The set reward probabilities pair per session were 0.1 Vs 0.4. The two left panels show the distribution of the block lengths and the difference in set reward probabilities. The two right panels show their correspondent reward and choice history effects. The correlations of the block lengths with the coefficients of reward and choices one trial back are reported as R on the plots with their correspondent significance labeled as *p* < 0.05 = *, *p* < 0.01 = **, *p* < 0.001 = * * *, and n.s. states for a non-significant result.

**Supplementary Figure 4:**
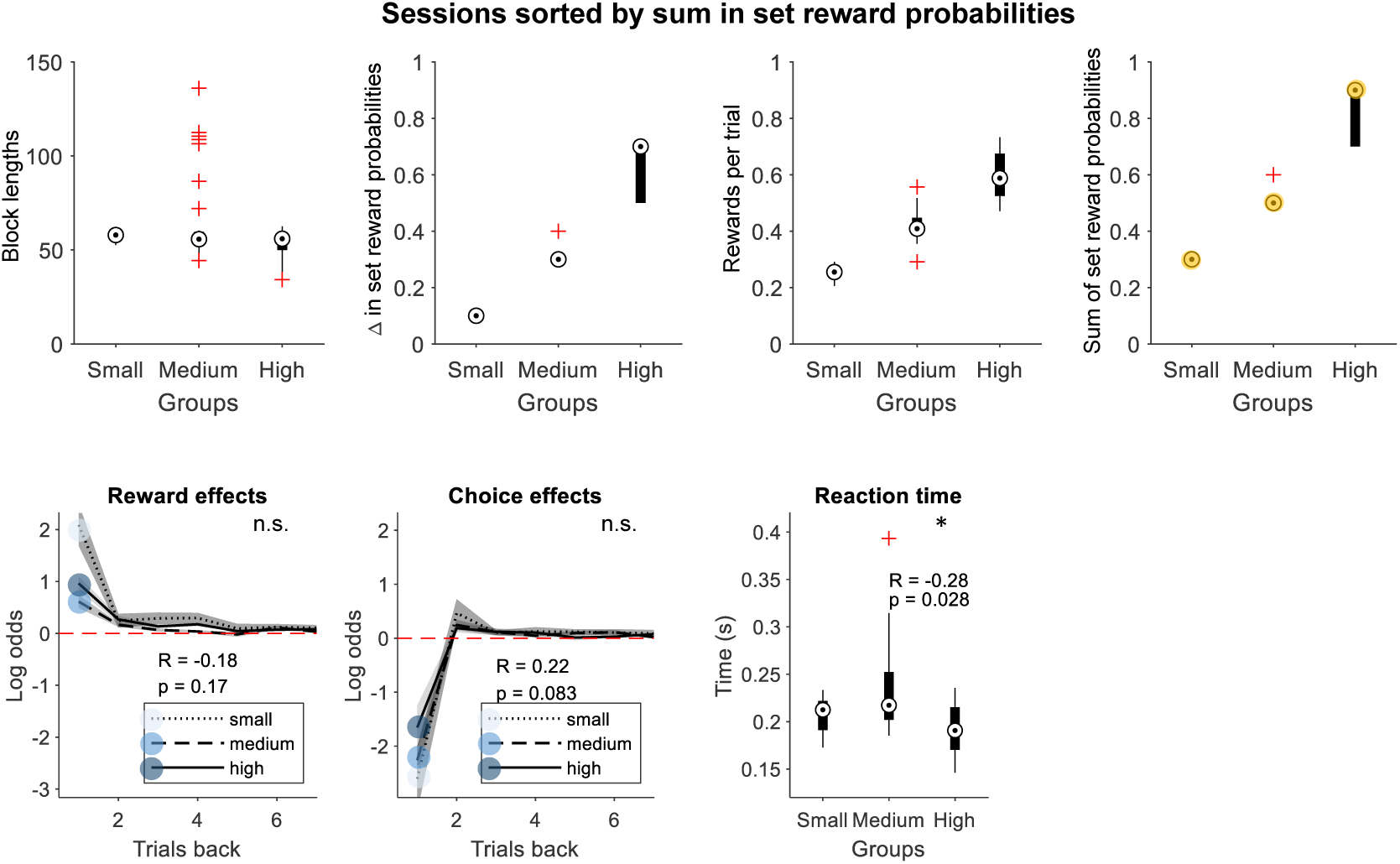
Increase in the total probability of reward delivery decreases the reaction time of mice. Sessions of four mice sorted in three groups (terciles) by the mean in set reward probabilities (62 sessions), or in other words, the mean set probability of rewards in a session. The set reward probabilities of the leaner side was kept to 0.1, while the richer side was set to 0.4, 0.5, 0.6 or 0.8 in a session. The upper panels show the distribution of the block lengths, the difference in set reward probabilities, the average number of rewards collected per trial and the mean of the set probabilities. The bottom panels show their correspondent reward history effects, choice history effects and reaction times. The correlations of the block lengths with the coefficients of reward and choices one trial back are reported as pearson correlation coefficient (r) on the plots with their correspondent significance labeled as *p* < 0.05 = *, *p* < 0.01 = ** and *p* < 0.001 = ***.

**Supplementary Figure 5:**
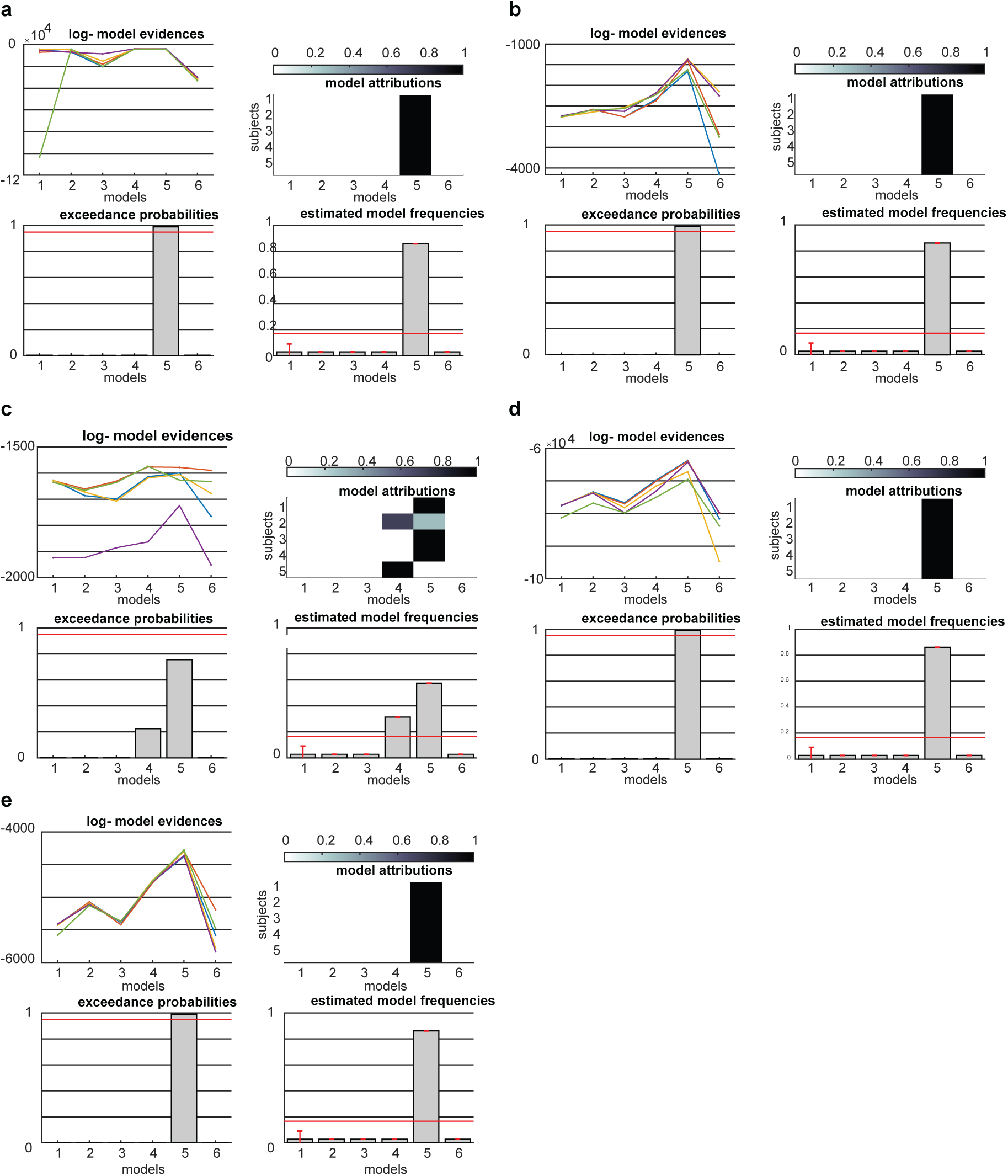
Bayesian Model selection. Numbers on x axis indicate different models. 1- Indirect model, 2 -F-Q, 3 - F-Q up, 4- F-Q W/C, 5 - Double trace, 6- Generalized linear model (a). MVT agent, (b) Oracle agent,(c) Human, (d) mice and (e) DT agent.

## Notes

### Competing Interest Statement

The authors have declared no competing interest.

